# Systematic Discovery of Novel Phosphoinositide-Binding Effectors in *Legionella* Reveals Conserved α-Helical Folds

**DOI:** 10.1101/2025.11.26.690877

**Authors:** Abigail E. Bolt, Abby E. Richardson, Colleen M. Pike, Rebecca R. Noll, Yanbao Yu, Joshua P. Neunuebel, Sylvain Le Marchand, Karl R. Schmitz, M. Ramona Neunuebel

## Abstract

Intracellular bacterial pathogens deploy effector proteins to hijack host membranes by targeting specific lipids. Phosphoinositides are key eukaryotic signaling molecules that govern membrane identity and dynamics, making them powerful levers for microbial interference. However, the bacterial repertoire of phosphoinositide-binding effectors remains largely undefined. In this study, we systematically identified such proteins using an expression library of 241 *Legionella pneumophila* effectors. We applied a combination of lipid bead pulldowns, mass spectrometry, colocalization with phosphoinositide biosensors, and protein-lipid overlay assays, leading to the validation of 18 new phosphoinositide-binding effectors. Structural predictions revealed that these proteins share compact alpha-helical folds distinct from canonical eukaryotic lipid-binding domains. Guided by this structural signature, we performed a computational screen across *L. pneumophila*, *Coxiella burnetii*, and *Burkholderia pseudomallei*. This analysis identified 15 additional effectors, which were experimentally validated as phosphoinositide binders. Altogether, we report 33 previously unrecognized bacterial phosphoinositide-binding effectors. While their lipid-binding profiles were diverse, many effectors showed a preference for compartments enriched in phosphatidylinositol 3-phosphate. These findings expand the known diversity of lipid-binding effectors produced by intracellular pathogens, identify new structural modules bacteria use for phosphoinositide recognition, and broaden our understanding of how pathogens exploit phosphoinositide signaling to manipulate host membranes.

**Importance:** Intracellular bacterial pathogens hijack host membrane trafficking by targeting phosphoinositides (PIPs), essential eukaryotic signaling lipids. Despite their importance in virulence, bacterial PIP-binding effectors remain largely uncharted because they lack recognizable sequence motifs. Using Legionella pneumophila as a model system, we systematically identified 30 novel PIP-binding effectors, doubling the known repertoire, and discovered that they employ compact α-helical folds structurally distinct from canonical eukaryotic PIP-binding domains. Remarkably, these α-helical modules are conserved in *Coxiella burnetii* and *Burkholderia pseudomallei*, revealing a convergent evolutionary strategy for PIP recognition across phylogenetically diverse pathogens. This work expands our understanding of bacterial lipid-targeting mechanisms and provides a structural blueprint for identifying virulence factors in emerging pathogens.

## Introduction

Intracellular bacterial pathogens have developed an extensive array of strategies to manipulate host cell functions and create environments that support their replication. A key aspect of this manipulation involves hijacking host cellular membranes, a process often mediated by bacterial effector proteins that are injected into the host cytosol via specialized secretion systems. One common tactic among diverse pathogens remodeling host membranes is targeting specific lipids. Among these, phosphoinositides (PIPs) have emerged as critical targets due to their central role in regulating membrane identity and dynamics^1,2^. Phosphatidylinositol (PI), the backbone of PIPs, consists of a membrane-anchored fatty acid tail and a myo-inositol headgroup that can be phosphorylated at the 3′, 4′, and 5′ positions to generate seven distinct PIP species^1^. Though they comprise a small fraction of total lipids, PIPs are distributed across specific membrane compartments where they regulate vesicle trafficking, cytoskeletal organization, signal transduction, and membrane dynamics^3,4^. Unraveling the molecular details of how pathogens target and manipulate PIPs is crucial to understanding how they gain control over host membranes, facilitating their entry, intracellular survival, replication, and evasion of immune defenses.

Bacterial effectors that bind PIPs have been identified in diverse intracellular pathogens, including *Shigella flexneri*, *Salmonella enterica*, *Mycobacterium tuberculosis*, *Listeria monocytogenes*, *Francisella tularensis, Coxiella burnetii*, and *Legionella pneumophila*^5–12^. By sensing specific PIPs, these effectors localize to defined membrane compartments where they influence vesicle identity, membrane curvature, and the recruitment of host machinery to support bacterial replication, highlighting PIP recognition as a core virulence strategy. Although PIP-targeting effectors play critical roles, identifying them remains a significant challenge. Unlike eukaryotic PIP-binding proteins, which contain conserved domains such as Pleckstrin Homology (PH), Fab1-YOTB-Vac1-EEA1 (FYVE), or Phox Homology (PX) domains^13^, bacterial effectors often lack recognizable motifs, making them hard to detect by sequence alone^14,15^. To overcome this, approaches that detect functional convergence beyond primary sequence are needed. With its large effector arsenal (>330) and robust experimental tools, *Legionella pneumophila* serves as an ideal model for such discovery efforts^16–18^.

*Legionella pneumophila (*hereafter *Legionella)* is a facultative intracellular pathogen that infects both environmental amoebae and human alveolar macrophages, causing the often-severe pneumonia known as Legionnaires’ disease^19–21^. Upon phagocytic uptake, *Legionella* resides within an early phagosome that it rapidly remodels into a replication-permissive compartment known as the *Legionella*-containing vacuole (LCV)^22,23^. Rather than following the canonical maturation pathway toward lysosomal degradation, this initial phagosome is extensively reprogrammed as *Legionella* diverts host vesicular traffic and alters membrane composition through the concerted action of numerous effector proteins secreted via the Dot/Icm type IV secretion system^24,25^. Early in infection, the LCV acquires endoplasmic reticulum (ER)-derived vesicles^26–28^ and becomes enriched in phosphatidylinositol 4-phosphate [PI(4)P]^29^, distinguishing it from conventional phagosomes. A subset of *Legionella* effectors plays a direct role in this membrane remodeling by targeting host PIP signaling. These effectors employ both lipid recognition and enzymatic activity, some mimicking host enzymes to generate specific PIP species at the LCV^11^, underscoring lipid manipulation as a central feature of *Legionella*’s intracellular strategy. Following initial PI(3)P accumulation on the LCV^29,30^, PI(3)P is converted to PI(3,4)P_2_^31^, and then shifts to PI(4)P which is gradually enriched on the vacuole membrane in an effector-dependent manner^29,32^. This PI(4)P-rich platform selectively recruits lipid-binding effectors. Notably, SidM (DrrA) and SidC, the first identified PIP-binding effectors, anchor to PI(4)P via distinct domains, P4M and P4C, respectively^33,34^. SidM functions as a Rab1 GEF, promoting recruitment and fusion with ER-derived vesicles, while SidC acts as an E3 ubiquitin ligase driving recruitment of ER-derived components^34–37^. The early discovery of PI(4)P-binding effectors, coupled with the PI(4)P enrichment on the LCV, initially suggested a dominant role for PI(4)P in effector targeting. However, many subsequently identified effectors were found to preferentially bind PI(3)P^38–42^, a lipid more commonly associated with early endosomes and autophagic membranes, revealing a broader and more nuanced role for PIP signaling in pathogenesis. This supports a dynamic model where PI(3)P- and PI(4)P-binding effectors function at distinct stages or subdomains to drive membrane remodeling and immune evasion.

In *Legionella*, several conserved PIP-binding modules have been identified, including the PI(4)P-binding P4M^33^ and P4C domains^34^, and the PI(3)P-specific LED modules^38^. Since many known *Legionella* PIP-binders do not harbor these modules, we hypothesized that more PIP-binding modules likely exist beyond those already identified. While sequence-based genomic analyses have been effective in identifying conserved domains, they are limited in their ability to detect structurally divergent or cryptic features. Investigating whether bacteria, like their eukaryotic hosts, harness a wider array of strategies to engage membrane lipids is a crucial next step in uncovering the full complexity of effector-membrane interactions.

To systematically identify PIP-binding effectors in *Legionella*, we employed a comprehensive multipronged approach combining biochemical and cellular assays. Our experimental workflow began with a high-throughput pulldown assay using lipid-coated beads and mass spectrometry to identify PIP-binding candidates from an expression library of 241 *Legionella* effectors. Candidates were then assessed by confocal imaging for subcellular distribution and co-localization with PI(3)P and PI(4)P biosensors, followed by protein-lipid overlay assays to probe direct binding. Through this approach, we revealed 18 novel PIP-binding effectors, most of which showed strong specificity for PI(3)P. Notably, based on predicted AlphaFold models, many of these effectors contained uncharacterized α-helical regions, which we experimentally confirmed as novel PIP-binding folds. Using this structural signature as a guide, we performed a structure-based bioinformatic screen against a database of *Legionella* effector domains using Flexible structure AlignmenT by Chaining Aligned fragment pairs allowing Twists (FATCAT), identifying additional candidate PIP-binding proteins. Experimental validation confirmed 12 of these candidates, bringing the total number of PIP-binding effectors identified in this study to 30 and substantially expanding the known arsenal of PIP-targeting proteins in *Legionella*. To investigate whether these newly identified PIP-binding folds are structurally conserved in other pathogens, we performed structure-based homology searches against effector repertoires from *Coxiella burnetii* and *Burkholderia pseudomallei*. This analysis revealed the presence of effectors with similar α-helical folds, and we validated PIP-binding activity in two *Coxiella* proteins and one *Burkholderia* effector. These results demonstrate PIP interaction through previously unrecognized, structurally conserved folds as a shared strategy among diverse intracellular pathogens. Together, these findings reveal a convergent mechanism by which bacterial effectors engage host lipid signaling and provide a framework for systematically identifying proteins that manipulate host membranes across diverse pathogens.

## Results and Discussion

### Optimization of Expression and Solubility Conditions for a *Legionella* Effector Library

Despite the identification of a growing number of PIP-binding effectors, only a handful share recognizable PIP-binding domains, and they show little to no resemblance to canonical eukaryotic PIP-binding modules. Whether additional, more broadly conserved structural features exist among PIP-binding effectors has remained an open question. Resolving this requires a large set of experimentally validated PIP-binding effectors for meaningful comparisons. Important advances have been made by prior studies using bioinformatic approaches to predict candidates, but methods relying on specific sequence or structural motifs may overlook effectors with unconventional features. To overcome this limitation, we conducted a comprehensive biochemical screen across a broad *Legionella* effector library to directly identify novel PIP-binding proteins.

To this end, we used an *Escherichia coli* BL21(DE3) Gateway expression library harboring 241 effectors with an N-terminal 6×His tag. Given that many *Legionella* effectors associate with membranes, and that membrane-associated proteins are often prone to poor solubility, we reasoned that optimizing expression and stability conditions would be critical to maximize our chances of obtaining sufficient, properly folded protein for downstream lipid-binding assays (**Supplementary Fig. S1A**). We first compared effector expression levels in Luria-Bertani (LB) medium and MagicMedia™ for four effectors: Lpg0696/Lem3, Lpg1950/RalF, Lpg2411/Lem24, and Lpg2464/SidM. Each showed markedly higher expression in MagicMedia™ compared to LB (**Supplementary Fig. S1B**). Based on these results, we selected MagicMedia™ for large-scale overproduction of the effector library to support subsequent screening assays.

Recombinant proteins expressed in *E. coli* often unfold upon lysis, leading to aggregation and loss of solubility ^43^. To improve recovery, we supplemented the lysis buffer with chemical chaperones. Testing trehalose and L-arginine using 6×His-Lpg2411/Lem24 as a representative, we found that both increased the amount of soluble protein compared to the standard buffer (**Supplementary Fig. S1C**). However, given L-arginine’s positive charge and the risk of nonspecific interactions with the negatively charged PIP-coated beads, we selected trehalose, a nonionic additive, for subsequent experiments. Proof-of-concept pulldown assays with Lpg2464/SidM, a known PI(4)P-binding effector, and Lpg1950/RalF, a non-binder ^33^, confirmed that trehalose preserved specific binding without introducing nonspecific interactions (**Supplementary Fig. S1D**). Having established optimized conditions for protein recovery, we next screened the effector library for PIP-binding activity using PIP-coated beads.

### Mass Spectrometry-Based Identification of Candidate PIP-Binding Effectors

To uncover novel PIP-binding effectors, we screened a 241-strain *Legionella* library using lipid-coated beads followed by mass spectrometry-based candidate identification (**Fig. 1**). To enable efficient screening of the *Legionella* effector library, we organized the effectors into 31 pools of 7–8 strains each, including one known PIP-binding effector per pool as internal positive controls (**Supplementary Data S1**). Each strain was cultured separately to maximize protein yield and prevent competition effects that could skew effector representation in pooled lysates. After growth, strains within each pool were combined and lysed. Whole-cell lysates were cleared by centrifugation to remove cellular debris, then incubated with control agarose, PI(3)P-coated, and PI(4)P-coated beads. A total of 93 eluates were collected and analyzed by mass spectrometry (MS) (**Fig. 1**), leading to the identification of 100 effectors; including 11 known PIP-binders, resulting in 89 new candidates (**Supplementary Data S1**).

**Figure 1.**
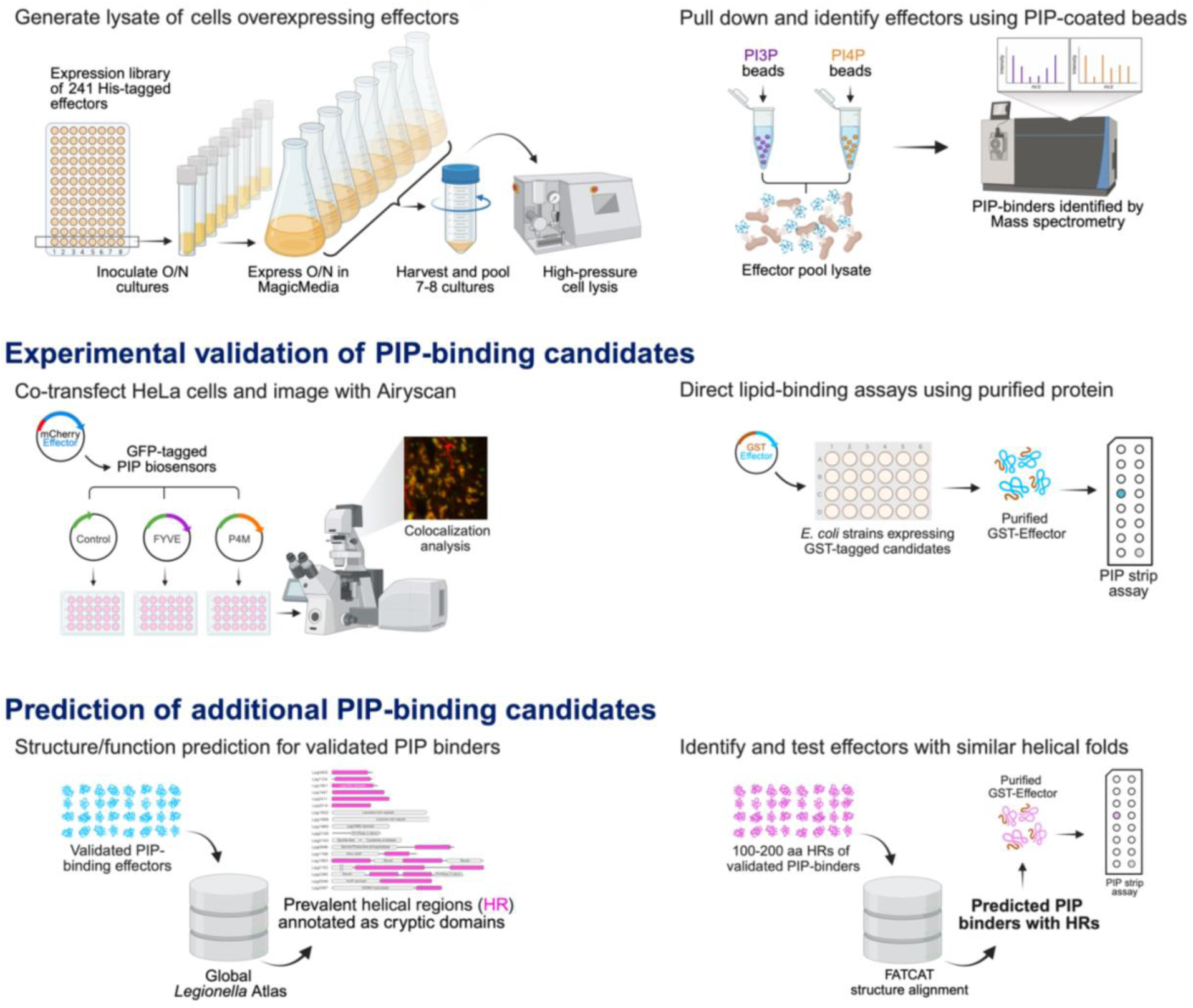
Workflow for discovery and validation of novel PIP-binding effectors. A multi-step pipeline was used to uncover novel PIP-binding effectors in *Legionella*. Lipid-bead pulldown followed by mass spectrometry identified candidate effectors. Co-localization with PIP biosensors in mammalian cells prioritized effectors with membrane targeting potential. Protein-lipid overlay assays biochemically validated direct PIP binding. AlphaFold-based structural analysis identified conserved α-helical regions among validated binders. Structure-guided prediction and FATCAT similarity clustering uncovered additional candidates, leading to the identification of new PIP-binding effectors.

Although control beads occasionally yielded similar or higher peptide counts compared to PI(3)P- or PI(4)P-coated beads (including for known PIP-binding effectors), the MS screen provided an effective starting point for candidate identification. Most hits did not display strong selectivity between PI(3)P and PI(4)P beads based on peptide counts. Given these observations and the large number of candidates recovered, we used the MS results as a foundation for a focused and systematic validation pipeline. We subsequently implemented two complementary approaches: co-localization studies with PIP-biosensors and protein-lipid overlay assays.

### Colocalization with PIP Biosensors Identifies High-Priority Candidate Effectors

To strengthen our candidate selection beyond MS-based identification, we next assessed whether effector localization correlated with PIP-enriched membranes in mammalian cells. We generated N-terminal mCherry fusion constructs for each of the 89 candidate effectors (**Supplementary Fig. S2**) and transiently co-transfected HeLa cells with plasmids encoding the mCherry-tagged effector and either GFP-2×FYVE or GFP-2×P4M, biosensors for PI(3)P and PI(4)P, respectively ^44^ (**Fig. 1**). Using confocal microscopy, we evaluated the extent of colocalization between each effector and the PIP biosensors. Of the 89 candidates, we successfully acquired images for 85; mCherry-Lpg0944/RavJ caused extensive cell rounding, and mCherry-Lpg1355/SidG, mCherry-Lpg1368/Lgt1, and mCherry-Lpg2244 were expressed at levels too low for detection. Of the 85 effectors analyzed, 25 showed overlap with the biosensors based on visual examination and we further quantified their extent of overlap using Mander’s Overlap Coefficient (MOC), where a value of 1 indicates complete overlap and 0 indicates random localization ^45^ (**Fig. 2A; Supplementary Fig. S3 & S4**). The majority of these effectors displayed preferential localization to PI(3)P-enriched compartments with 15 strongly colocalizing effectors (MOC > 0.6) and 5 moderately colocalizing effectors (MOC 0.6 > 0.4). Although Lpg1148 /LupA and Lpg1689 showed weak overall colocalization with FYVE (MOC ∼0.3), the values were statistically greater than those observed with P4M, suggesting a low-level but specific enrichment at PI(3)P-enriched compartments. Lpg1958/LegL5 was the only effector to preferentially localize to PI(4)P-enriched structures. Lastly, Lpg1963/PieA and Lpg2153/SdeC did not colocalize with either marker, however, they displayed adjacent localization with GFP-2×FYVE. Together, this colocalization analysis yielded a subset of 25 effectors and provided independent, focused evidence supporting the notion that these candidates may bind PIPs.

**Figure 2.**
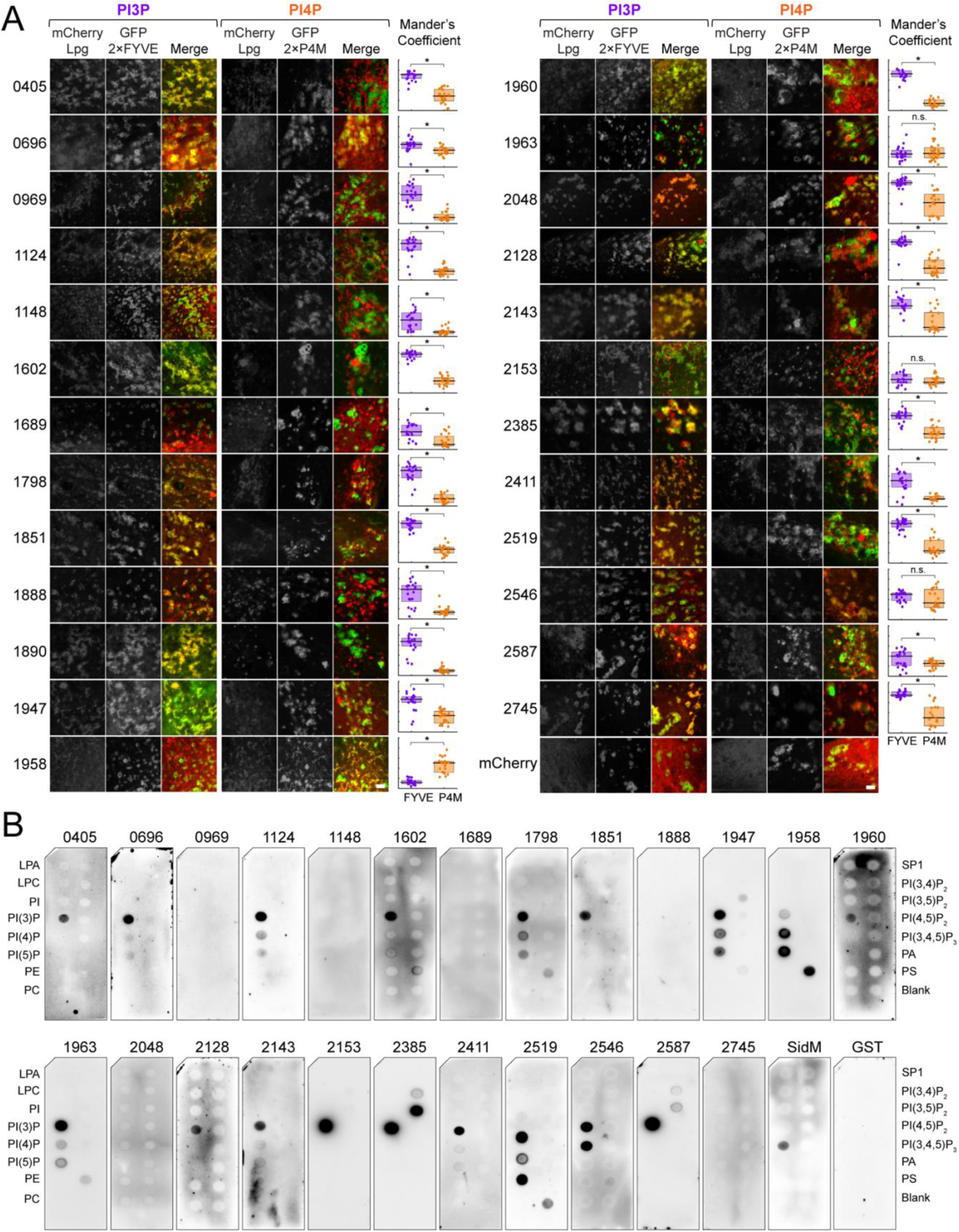
Identification and experimental validation of candidate PIP-binding *Legionella* effectors using colocalization and lipid binding assays. (A) Confocal microscopy insets of HeLa cells co-transfected with mCherry-tagged candidate effectors and either GFP-2×FYVE or GFP-2×P4M PIP biosensors (full size images in **Supplementary Fig. S3**). Scale bars, 2 μm. Plots showing Mander’s Overlap Coefficients (the fraction of red pixels that overlap with green pixels) with 21-26 independent data points per condition represented by individual dots. Median (black horizontal line) and interquartile range (purple box for FYVE and orange box for P4M). Y-axis ranges from 0-1 with ticks at every 0.2. Wilcoxon rank sum test, * denotes p < 0.05. (B) Protein-lipid overlay assays showing that 18 GST-tagged effectors bound one or more PIPs. Nitrocellulose membranes pre-spotted with 100 pmol of each lipid were incubated with purified effectors. Bound proteins were detected using an anti-GST-HRP conjugated antibody. Lipid abbreviations: LPA, lysophosphatidic acid; LPC, lysophosphocholine; PI, phosphatidylinositol; PE, phosphatidylethanolamine; PC, phosphatidylcholine; S1P, sphingosine-1-phosphate; P, phosphate; P2, bisphosphate; P3, triphosphate; PA, phosphatidic acid; PS, phosphatidylserine.

### Biochemical Validation Expands the Repertoire of *Legionella* PIP-Binding Effectors

To determine whether candidate effectors directly bound PIPs, we generated N-terminal GST fusion constructs for each of the 25 effectors. We expressed and purified 24 fusions (all except Lpg1890/LegLC8) and performed *in vitro* protein-lipid overlay assays (**Supplementary Fig. S5**). Purified effectors were incubated with nitrocellulose membranes pre-spotted with various lipids, including the seven major PIP species. A total of 18 effectors exhibited binding to one or more lipids (**Fig. 2B**). Strikingly, all but one effector (Lpg1958/LegL5) bound PI(3)P, either exclusively or alongside other lipids. Lpg1947/Lem16 bound PI(3)P and, to a lesser degree, PI(4)P and PI(5)P; Lpg1958/LegL5 bound PI(4)P, PI(5)P, and phosphatidylserine (PS). Lpg2385 bound PI(3)P and PI(3,5)P_2_, Lpg2519 bound PI(3)P and PI(5)P, and Lpg2546 bound PI(3)P and PI(4)P. The remaining effectors - Lpg0405, Lpg0696/Lem3, Lpg1602/LegL2, Lpg1798/MavU, Lpg1851/Lem14, Lpg1960/LirA, Lpg1963/PieA, Lpg2128, Lpg2143, Lpg2153/SdeC, and Lpg2587 - exhibited specific binding to PI(3)P. These findings substantially expand the known repertoire of *Legionella* PIP-binding effectors, increasing their number from 24 to 42 and providing new insights into the potential roles of previously uncharacterized proteins.

Several of the newly identified PIP-binding effectors have established roles in host manipulation, and PIP-binding adds a new dimension to their functional characterization (**Supplementary Table S1**). Lpg0696/Lem3, known to dephosphocholinate Rab1 and reverse AnkX-mediated modifications ^46^, now appears capable of directly associating with PIP-containing membranes. Lpg1851/Lem14, which acts in synergy with the PI3-phosphatase Lpg0130/SidP ^30,47^ exhibited PI(3)P binding, suggesting it co-targets PI(3)P-enriched structures generated by Lpg0130/SidP. Lpg1963/PieA, which localizes to the LCV and alters lysosomal morphology ^48^, may leverage its PI(3)P-binding ability to modulate endosomal dynamics. Lpg2153/SdeC, a SidE family member that modifies ER and Golgi proteins via non-canonical phosphoribosyl-ubiquitination ^49–51^, also bound PI(3)P, suggesting that PIP-binding may underlie its LCV association.

The remaining newly identified binders are functionally uncharacterized, and their ability to associate with PIPs provides an important first step toward defining their roles. For the six effectors that colocalized with PIP biosensors but did not show binding on lipid strips (Lpg0969/RavK, Lpg1148/LupA, Lpg1689, Lpg1888/LpdA, Lpg2048, and Lpg2745), several factors could explain the discrepancy. Localization may require membrane curvature not captured in the overlay assay, involve low-affinity interactions that are disrupted by the assay conditions, or is driven by association with another membrane component. Alternatively, binding could depend on another effector, complex formation, or post-translational modifications absent in the recombinant system. Having expanded the repertoire of PIP-binding effectors, we next asked whether these newly identified effectors shared any common structural features with each other or with previously characterized PIP-binding domains.

### Compact α-Helical Folds Mediate PIP-Binding by Many *Legionella* Effectors

To explore conserved structural features among newly identified PIP-binding effectors, we used the *Legionella* effector database developed by Patel et al. ^15^ which provides a comprehensive structural annotation of the 368 Dot/Icm-translocated effectors, integrating AlphaFold-predicted models with curated domain predictions. The majority of newly identified PIP-binders (13 of 18) contained uncharacterized α-helical regions or bundles, typically composed of 4 to 6 helices, in addition to various predicted or known functional domains (**Fig. 3A**). Notably, similar small α-helical folds were also present in 19 of 24 previously characterized PIP-binders, corresponding to their lipid-binding domains. Furthermore, the structurally defined PIP-binding modules - P4M from Lpg2464/SidM, P4C from Lpg2511/SidC, and the C-terminal domain of Lpg1683/RavZ-are also compact α-helical bundles comprising 4 to 6 helices (**Fig. 3A**) ^39,52,53^. Seven newly identified PIP-binders contained uncharacterized α-helical bundles alongside other functional domains, while another six were small (<275 amino acids) entirely α-helical proteins of unknown function (**Fig. 3A**). Lpg1851/Lem14 and Lpg1960/LirA, have been structurally characterized and also display predominantly α-helical architectures^47,54^. Interestingly, two additional PIP-binders, Lpg1958/LegL5 and Lpg1602/LegL2, contained Leucine-rich repeats (LRRs) rather than compact helical bundles. This mirrors previous findings that Lpg1660/LegL3, another LRR-containing effector, can bind PIPs^55^ and although we were unable to purify candidate Lpg1890/LegLC8, it also belongs to the LRR family. These motifs are typically involved in protein-protein interactions^56^ and have not been previously implicated in direct lipid binding. Their association with PIP-binding in our dataset suggests an unexpected functional versatility of the LRR scaffold in mediating lipid engagement.

**Figure 3.**
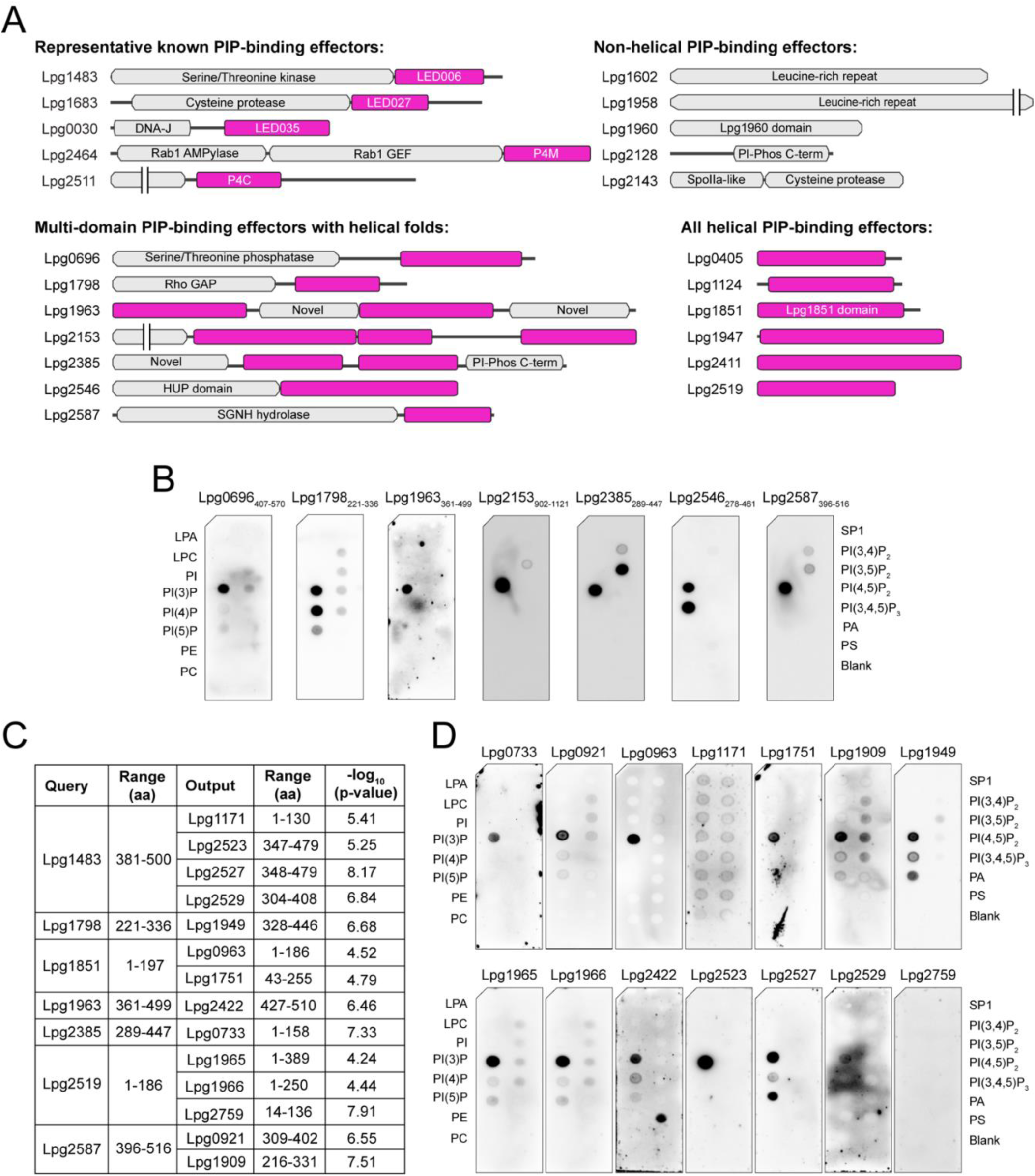
Alpha-helical folds mediate PIP-binding by *Legionella* effectors and can serve as signature features to identify additional effectors with PIP-binding activity. (A) Domain architecture of newly identified PIP-binding effectors predicted using AlphaFold models and curated domain annotations^15^. Uncharacterized α-helical regions identified in PIP-binders are highlighted in magenta. Effectors are grouped by domain organization, with multi-domain proteins shown separately from small, fully α-helical proteins (<275 amino acids) and LRR-containing effectors. (B) Protein-lipid overlay assays showing that GST-tagged truncation variants comprising selected αα-helical regions preserved the PIP-binding profiles of their corresponding full-length proteins. Nitrocellulose membranes pre-spotted with 100 pmol of each lipid were incubated with purified truncation variants. Bound proteins were detected using an anti-GST-HRP conjugated antibody. (C) Table summarizing a sample of hits from FATCAT analysis. (D) Protein-lipid overlay assays showing that 18 GST-tagged effectors bound one or more PIPs. Nitrocellulose membranes pre-spotted with 100 pmol of each lipid were incubated with purified effectors. Bound proteins were detected using an anti-GST-HRP conjugated antibody. Lipid abbreviations: LPA, lysophosphatidic acid; LPC, lysophosphocholine; PI, phosphatidylinositol; PE, phosphatidylethanolamine; PC, phosphatidylcholine; S1P, sphingosine-1-phosphate; P, phosphate; P2, bisphosphate; P3, triphosphate; PA, phosphatidic acid; PS, phosphatidylserine.

Given the prevalence of uncharacterized α-helical regions among both newly identified and previously known PIP-binders, we hypothesized that these folds mediate PIP-binding. To test this hypothesis, we expressed and purified GST-fusion truncation constructs spanning the uncharacterized helical regions of seven multi-domain effectors: Lpg0696/Lem3_407-570_, Lpg1798/MavU_221-336_, Lpg1963/PieA_361-499_, Lpg2153/SdeC_902-1121,_ Lpg2385_289-447_, Lpg2546_278-461_, and Lpg2587_391-516_ (**Supplementary Fig. S5**). For effectors with multiple helical regions (Lpg1963/PieA, Lpg2385, and Lpg2153/SdeC), we selected the region with the highest similarity to known PIP-binding folds based on structural analyses. Truncations were not generated for fully α-helical proteins under 275 amino acids, which may require the intact fold for structural stability (Lpg0405, Lpg1124, Lpg1851/Lem14, Lpg1947/Lem16, Lpg1960/LirA, Lpg2411/Lem24, and Lpg2519), or for LRR-containing effectors (Lpg1602/LegL2, Lpg1958/LegL5), where binding may involve extended surfaces across multiple repeats rather than localized domains. We next tested whether truncated variants comprising the α-helical regions preserved PIP-binding activity seen in the full-length proteins. Protein-lipid overlay assays showed that all tested truncation variants preserved PIP-binding profiles comparable to their full-length counterparts (**Fig. 3B**), confirming these α-helical regions as functional PIP-binding domains. Collectively, these results demonstrate that *Legionella* effectors predominantly engage PIPs through compact α-helical folds. This strategy appears widespread among both newly identified and previously characterized PIP-binders, highlighting a common structural principle underlying effector-mediated subversion of host membranes.

### Conserved Helical Folds Predict PIP-Binding Across *Legionella* Effectors

Despite some sequence homology within specific PIP-binding groups, the complete set of *Legionella* PIP-binding effectors lacks any universally conserved sequence motifs. However, given that many PIP-binding effectors contain α-helical folds, we hypothesized that structural homology could serve as a predictive tool for identifying additional PIP-binding candidates. To test this, we performed structural similarity searches against a curated database of 660 *Legionella* effector domains^15^ (**Fig. 1**). FATCAT’s tolerance for local flexibility makes it well suited to detect conserved folds among effectors with substantial sequence divergence^57^. Validated PIP-binding regions were used as queries when available; otherwise, full-length proteins were used (e.g., Lpg1660/LegL3, Lpg2222/LpnE). For newly identified binders, we used either confirmed binding domains or full-length sequences for small α-helical proteins and LRR-containing effectors. Structural comparisons yielded *p*-values quantifying the likelihood that alignments occurred by chance, with lower values indicating stronger structural similarity. These results were visualized as a clustered similarity matrix by plotting –log_10_(p-value) for each alignment across the effector dataset (**Supplementary Fig. S6**).

Several regions of high structural similarity emerged from the matrix, corresponding to regions where multiple effectors shared strong resemblance to the input PIP-binding domains. Within these areas, we identified clusters of effectors defined by the presence of shared structural features including the P4M domain, the LED035 domain, LRRs, as well as a larger cluster spanning both effectors with the LED006 domain and those with the P4C domain (**Supplementary Fig. S6**). We also observed that this cluster included helical regions from several effectors that had not been previously evaluated for PIP-binding. Given the prevalence of significant structural similarity to known PIP-binders, we selected a group of 14 effectors from this area with varying degrees of resemblance to several different PIP-binding effectors (**Fig. 3C**). GST-fusion constructs of Lpg1171, Lpg2523/Lem26, Lpg2527/LnaB, Lpg2529/Lem27, Lpg1949/Lem17, Lpg0963, Lpg1751, Lpg2422/Lem25, Lpg0733/RavH, Lpg1965/PieC, Lpg1966/PieD, Lpg2759, Lpg0921/MavT, and Lpg1909/DenR were generated and tested for PIP-binding using protein-lipid overlay assays. These revealed all but Lpg1171 and Lpg2759 bound PIPs, again with a strong preference for PI(3)P, except for Lpg2422/Lem25 and Lpg2527/LnaB which exhibited additional binding to phosphatidylserine (PS) and PI(5)P, respectively. We note that several of these newly validated PIP-binders (e.g., Lpg0733/RavH, Lpg1751, Lpg1949/Lem17, Lpg1965/PieC, Lpg1966/PieD, Lpg2422/Lem25, Lpg2529/Lem27, Lpg2759) were originally identified in the PIP-bead pulldown screen but were not initially prioritized for further analysis due to cytosolic localization or lack of co-localization with PIP biosensors (**Supplementary Fig. S4**). Consequently, subcellular localization alone is not sufficient to predict PIP-binding capacity, highlighting the importance of direct biochemical validation.

Among the newly identified PIP-binders, several effectors including Lpg1909/DenR, Lpg1949/Lem17, Lpg2527/LnaB, and Lpg2529/Lem27 have been previously characterized (**Supplementary Table S1**). Notably, a subset of these effectors is located within genomic regions associated with horizontal gene transfer and domain shuffling^58,59^. For example, a previously defined region of genomic plasticity spanning Lpg1963/PieA to Lpg1976/LegG1^48^ includes three PI(3)P-binding effectors identified here: Lpg1963/PieA, Lpg1965/PieC, and Lpg1966/PieD. Their proximity suggests possible co-acquisition and functional coordination. Similarly, neighboring effectors Lpg2523/Lem26, Lpg2527/LnaB, and Lpg2529/Lem27 were all validated as PIP-binders, reinforcing the idea that *Legionella* may organize lipid-targeting effectors into genomic clusters to facilitate coordinated manipulation of host membranes. Additionally, Lpg0963 and Lpg1751 share the same fold as Lpg1851/Lem14^15^, supporting this domain mediates PI(3)P recognition. Collectively, this structure-guided approach led to the validation of 12 additional PIP-binding effectors, bringing the total identified in this study to 30 and highlighting that PIP targeting is more pervasive within the *Legionella* effector repertoire than previously recognized.

### Structural Classification of PIP-binding Helical Domains Reveals Common Folds

Distinct clusters and varying similarity scores suggested that PIP-binding effectors belong to discrete structural families (**Supplementary Fig. S6**). To systematically evaluate these relationships, we performed an all-against-all structural comparison using the DALI server, we analyzed the same structures used to generate the similarity matrix as well as helical regions from the 12 validated PIP-binders discovered through FATCAT analysis. The human FYVE domain from the EEA1 protein was included as an outgroup^60^. The resulting dendrogram grouped known PIP-binding domains as expected, positioning established domains such as P4M, P4C, LED006, LED035, and LED027 into distinct clusters (**Fig. 4**). Grouping was primarily driven by helix number and spatial organization. To further explore these relationships, we examined AlphaFold-predicted structures from each group to identify common fold features and potential mechanisms of PIP engagement.

**Figure 4.**
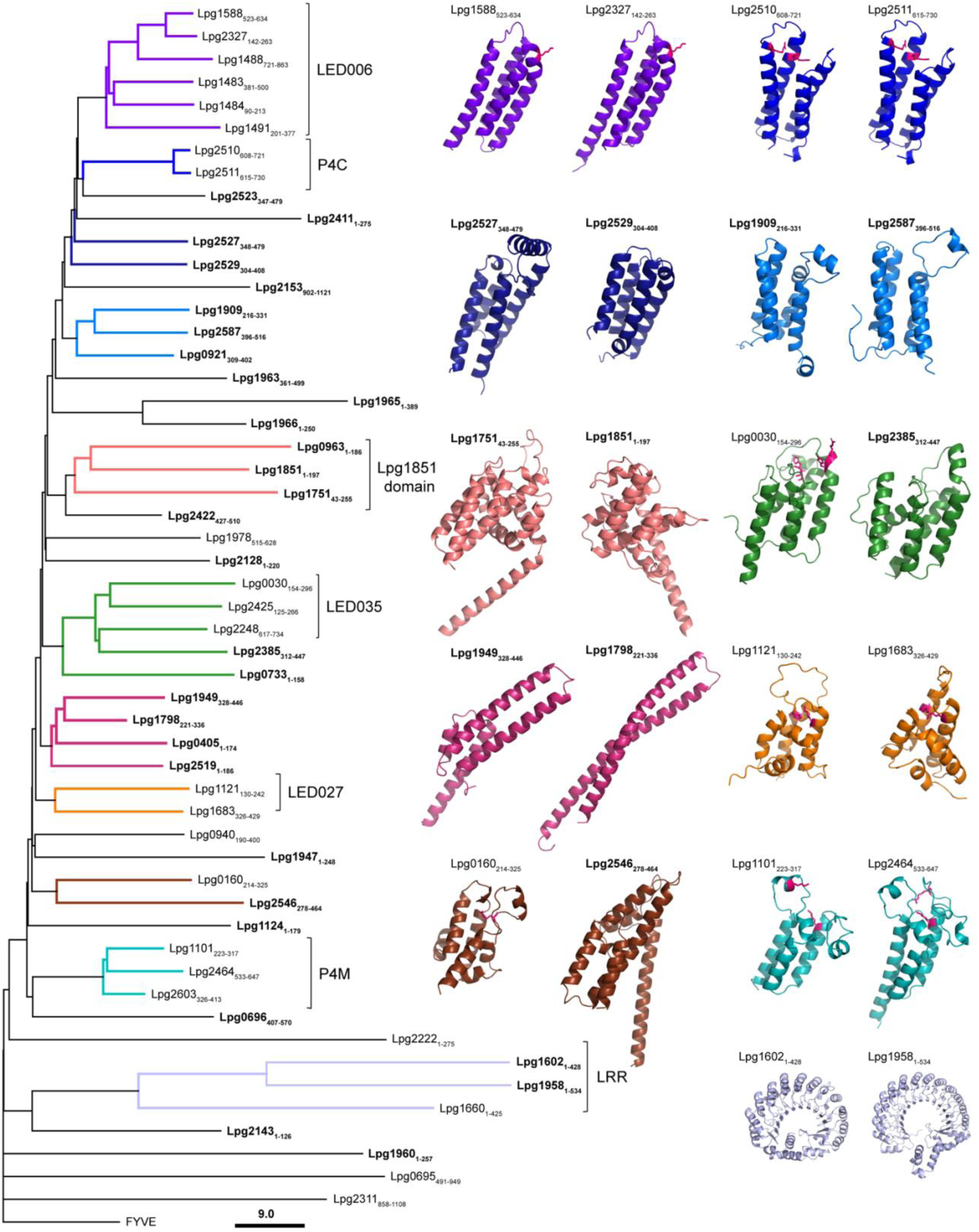
Hierarchical Clustering Reveals Distinct Helical Fold Families in *Legionella* PIP-Binding Effectors. All-against-all dendrogram of AlphaFold-predicted *Legionella* PIP-binding domains generated by DALI. Effectors with similar helical folds cluster together, revealing both known and novel fold families. Branch lengths reflect DALI Z-score–based structural dissimilarity, and node labels indicate each effector plus the aligned residue range. Branch colors define structural clusters by helix architecture: purple (LED006), blue (P4C), navy/marine (four-helix bundles), salmon (Lpg1851-like), green (five-to six-helix bundles, including LED035), berry (elongated 2–3-helix folds), orange (five-helix LED027), brown (seven-helix bundles), teal (P4M), and light blue (LRR). Labels in bold are PIP-binders discovered in this study. Experimentally validated PIP-binding residues are shown as pink sticks.

A structural comparison of groups reveals similarities that are not evident from sequence alone. For example, LED006- and P4C-containing proteins form separate but adjacent groups on the tree (**Fig. 4**) and are located within a larger cluster that includes newly identified PIP-binders (Lpg2523/Lem26, Lpg2411/Lem24, Lpg2527/LnaB, Lpg2529/Lem27, Lpg2153/SdeC, Lpg1909/DenR, Lpg2587, Lpg0921/MavT, and Lpg1963/PieA; **Fig. 4**). Although there is no notable sequence homology between groups, AlphaFold predictions all share a common core 4-helix bundle architecture. Positively charged residues that have been experimentally validated as critical for PIP-binding for LED006^38^ and P4C^52^ are located at one end of the helical bundles (**Fig.4, depicted as pink sticks**) near the α1-α2 and α3-α4 junctions and ideally positioned for lipid interaction. While this spatial arrangement was previously known for P4C, our analysis suggests that LED006 domains and other 4-helix bundle PIP-binders utilize an analogous mode of membrane engagement.

In a neighboring cluster, Lpg0733/RavH and Lpg2385, clustered with LED035 domain effectors despite lacking the LED035 motif, indicating a shared fold. Their AlphaFold models revealed a five-helix fold, sometimes with a sixth smaller helix nestled within a loop region. Like the LED006 and P4C domain proteins, the critical binding residues (**Fig.4, depicted as pink sticks**) in LED035-containing effectors were found within the loop region^38^. Similarly, LED027 domain-containing effectors formed five-helix bundles, although the helices were arranged differently from those in the LED035 group, suggesting the LED027 domain represents a distinct structural subclass. (**Fig. 4**). A separate group was formed by newly identified PIP-binders Lpg1949/Lem17, Lpg1798/MavU, Lpg0405, and Lpg2519. Unlike the more compact bundles described above, these structures were elongated and comprised of fewer helices, 2-3, suggesting a potential alternative mode of PIP engagement. Additional outliers included Lpg0160/RavD and Lpg2546, whose helical domains each consisted of seven α-helices, however, differences in helix length and packing resulted in structures that were not closely alike and distinct from other clusters (**Fig. 4**).

Another group included effectors sharing the Lpg1851-like fold as expected. These proteins were fully α-helical and contained a greater number of helices, but with orientations distinct from the four-helix bundles. Lastly, the previously described P4M domain of a six-helix fold composed of three long and three short helices formed a distinct cluster. As observed across most of the structural classes, the experimentally validated residues were consistently localized to loop regions and are shown in pink^32^ (**Fig. 4, teal**). Collectively, these findings demonstrate that variations in helix number and arrangement drive the structural clustering of PIP-binding effectors. Notably, many of the newly identified PIP-binding folds clustered with LED006 and LED035 domain-containing proteins, which were originally defined by sequence homology^38^. In addition to these established groups, our structure-guided analysis uncovered new PIP-binding folds sharing similar helical topologies. Despite their structural diversity, these folds converge functionally in their ability to engage PIPs.

### Structure-Based Discovery of Novel PIP-Binding Effectors in *Coxiella* and *Burkholderia*

Building on our structure-guided discovery of *Legionella* PIP-binding effectors, we asked whether the same strategy could reveal lipid-targeting proteins in other intracellular pathogens. We focused on *Coxiella burnetii* and *Burkholderia pseudomallei* due to their contrasting intracellular lifestyles; *Coxiella* thrives within an acidified lysosome-like vacuole^61^, whereas *Burkholderia* rapidly escapes into the host cytosol^62^. Despite these fundamental differences, both must subvert host membrane processes to survive. First, we performed FATCAT all-against-all alignments between ∼150 *Coxiella burnetii* Nine Mile RSA439 effectors fragments (200-residue AlphaFold segments) and the validated *Legionella* PIP-binding folds. The resulting similarity matrix highlighted a distinct region of high scoring alignments, from which we selected the top seven candidates, including CBU1677, CBU0041/CirA, and the known PI(3)P binder CBU0021/CvpB^10^ (**Fig. 5B**). Structural models of these three proteins revealed compact α-helical bundles closely matching the *Legionella* four-helix fold, with CBU1677 featuring an additional short fifth helix (**Fig. 5C**). We then generated GST-tagged constructs of full-length CBU1677 and the 1–200 region of CBU0041/CirA and confirmed their lipid specificity by protein-lipid overlay: CBU1677 bound both PI(3)P and PI(4)P, whereas CBU0041/CirA bound selectively to PI(3)P (**Supplementary Fig. S5; Fig. 5C**). Four other high-scoring hits displayed helical architectures and remain to be experimentally validated. Extending this workflow to *Burkholderia*, we identified the Type III effector BopA^63^: its 350–512 segment forms a four-helix bundle and binds specifically to PI(5)P (**Fig. 5D**), marking the first bacterial effector with a PI(5)P preference. Together, these findings indicate that the helical PIP-binding folds defined in *Legionella* are conserved among distinct intracellular pathogens and represent a shared virulence strategy among bacterial effectors that target host membranes. Additionally, they underscore the power of structure-based homology searches to uncover lipid-targeting effectors beyond the limits of sequence similarity.

**Figure 5.**
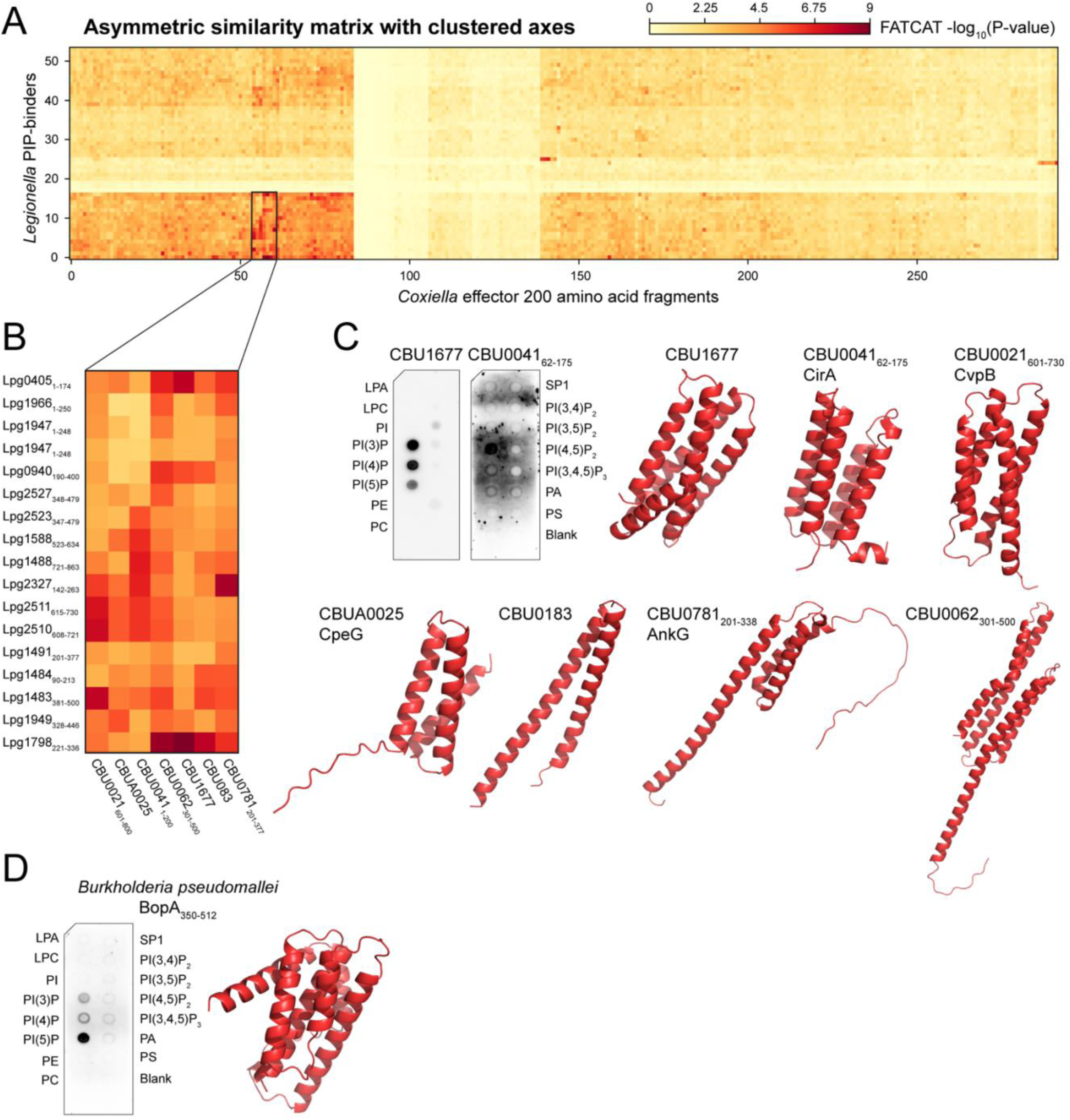
Conserved helical folds confer PIP-binding activity to effectors from *Coxiella burnetii* and *Burkholderia pseudomallei*. (A) A matrix was generated using the −log(P-value) from the comparison of PIP-binding effector domain structures (or full-length protein when the domain was unknown) to 200 amino acid AlphaFold fragments of *Coxiella* effectors by FATCAT. Orange to dark red represent statistically significant hits. Axes are clustered by Ward’s method. A box is drawn around an area of interest. (B) Magnified area of interest containing *Coxiella* effector hits with the largest −log(P-value) compared to numerous *Legionella* PIP-binding effectors across different fold groupings. (C) Protein-lipid overlay assays with purified GST-tagged effectors from *Coxiella,* CBU1677 and CBU0041_1-200_. Nitrocellulose membranes pre-spotted with 100 pmol of each lipid were incubated with GST fusion proteins, and bound proteins were detected using an anti-GST-HRP conjugated antibody. AlphaFold predicted structures for CBU1677, CBU0041_62-175_, CBU0021_601-730_, CBUA0025, CBU0183, CBU0781_201-338_, sand CBU0062_301-500_ depicting helical folds with similarity to *Legionella* PIP-binding effectors. Disordered regions were removed from CBU0041 (spanning residues 1-61 and 176-200) and CBU0021 (spanning residues 731-800). (D) Protein-lipid overlay assay with purified GST-tagged BopA_350-512_ from *Burkholderia*. Nitrocellulose membranes pre-spotted with 100 pmol of each lipid were incubated with GST fusion protein, and bound proteins were detected using an anti-GST-HRP conjugated antibody. AlphaFold predicted structure for BopA_350-512_. Lipid abbreviations: LPA, lysophosphatidic acid; LPC, lysophosphocholine; PI, phosphatidylinositol; PE, phosphatidylethanolamine; PC, phosphatidylcholine; S1P, sphingosine-1-phosphate; P, phosphate; P2, bisphosphate; P3, triphosphate; PA, phosphatidic acid; PS, phosphatidylserine.

In conclusion, we demonstrate that structurally homologous but topologically variable α-helical folds confer PIP-binding activity to a broad set of *Legionella* effectors, revealing a novel and prevalent mechanism of host membrane targeting. We identified 30 previously unrecognized *Legionella* PIP-binding effectors, bringing the total to 54, which accounts for a remarkable 20% of its known effector arsenal dedicated to co-opting host PIPs. This finding complements a growing body of work emphasizing the role of lipid targeting in bacterial pathogenesis and reveals that lipid-binding can be mediated by bacterial-specific PIP-binding modules with no homology to canonical eukaryotic PIP-binding motifs. Despite identifying numerous new PIP binders, our screen likely captures a subset of the full repertoire, and additional effectors, especially those with similar helical folds, may yet be uncovered. A striking number of effectors identified here and previously, bind PI(3)P, a degradative signpost that is primarily found on the compartments *Legionella* is avoiding in the cell. Although it remains unclear how each individual PI(3)P-binding effector is spatially and temporally regulated, their abundance indicates that *Legionella* heavily invests in targeting this lipid. Extending our analysis beyond *Legionella*, we identified homologous helical PIP-binding modules in *Coxiella burnetii* and *Burkholderia pseudomallei*, two pathogens with distinct intracellular lifestyles and survival strategies. Their shared use of structurally conserved, noncanonical lipid-binding folds underscores a unifying virulence mechanism through which diverse bacteria exploit host membrane systems to promote infection.

## Materials and Methods

### Strains, media, and reagents

All the strains used in this study are listed in **Supplementary Table S2**. HeLa cells (ATCC, CCL-2) were cultured at 37°C with 5% CO2 in RPMI 1640 supplemented with 2mM L-glutamine and 10% FBS. Antibodies were purchased from ProteinTech (HRP-conjugated 6*His, His-Tag mouse monoclonal, HRP-66005), Sigma (mouse monoclonal anti-β-actin, A2228), and ThermoFisher (mouse monoclonal anti-GST HRP-conjugated, MA4-004-HRP and rabbit polyclonal anti-mCherry, PA5-34974). MagicMedia™ was purchased from ThermoFisher (K6810).

### Construction of expression clones

Bacterial strains, plasmids, and oligonucleotides used in this study are listed in **Supplementary Table S3**. *E. coli* BL21 (DE3) expression library was constructed from a GC5 pDEST17 library of *L. pneumophila* effector proteins, a kind gift from Matthias Machner (National Institutes of Health). The pDONR221 effector constructs used in this study were obtained from a pDONR221 effector library, also gifted by Machner. GST and mCherry-fusion constructs of full-length and fragment effectors were generated by Gateway™ Cloning technology using the destination vectors pDEST™15 (ThermoFisher Scientific 11802014) and 362 pCS Cherry DEST, a kind gift from Nathan Lawson (Addgene plasmid #13075)^64^.

### Recombinant protein expression and purification

*E. coli* BL21 (DE3) or BL21 (DE3) pLysS stains carrying vectors encoding 6×His or GST-fusion proteins were grown in LB or MagicMedia™. First, starter cultures were grown overnight at 37°C. The next day, LB or MagicMedia™ was inoculated at 1:40 dilution from the starter culture and grown at 37°C until an OD600 ∼0.6 was reached. For LB cultures, 0.5 mM isopropyl 1-thio-β-D-galactopyranoside (IPTG, Sigma) was added and 6×His or GST-fusion proteins were produced at 18°C overnight.

GST-fusion proteins were purified using either glutathione 4B Sepharose™ (Cytiva) or MagneGST purification kit (Promega). For glutathione purification, *E. coli* BL21 (DE3) pLysS cells producing GST fusion effectors were harvested and resuspended in phosphate buffered saline (PBS; 11.9 mM phosphate, 137 mM sodium chloride, 2.7 mM potassium chloride) supplemented with 1 mM βME, 1 mM MgCl2, and 750 mM trehalose (PBS-MM + Trehalose) followed by lysis with LV1 Microfluidizer (Microfluidics). Cell lysate was centrifuged at 24,000 × g for 30 min and the supernatant was incubated with pre-equilibrated Sepharose 4B for 2 h at 4°C with end over end rotation. The resin was washed three times with PBS-MM and proteins were eluted in 50 mM Tris-HCl (pH 8) containing 10-50 mM reduced glutathione.

For MagneGST purification, GST-fusion effectors were purified according to manufacturer instructions (Promega). Briefly, *E. coli* BL21(DE3) pLysS cells producing GST fusion effectors were harvested and resuspended in MagneGST lysis buffer. Cells were incubated in the lysis buffer for 30 min at 4°C. Cell lysate was incubated with pre-equilibrated magnetic GST beads for 1 h at 4°C. The magnetic beads were washed three times with MagneGST wash/bind buffer and proteins were eluted in 50 mM Tris-HCl (pH 8) containing 50 mM reduced glutathione. After purification, glutathione was removed using a desalting Zeba column according to manufacturer instructions (ThermoFisher Scientific).

### PIP-bead pulldowns and mass spectrometry

*E. coli* BL21(DE3) cells expressing 6×His-tagged effectors were lysed in PBS-MM + Trehalose (pH 7.5) using a LV1 Microfluidizer (Microfluidics) and spun at 24,000 × g for 30 min. Of this lysate, 250μL was added to control agarose, PI(3)P-, and PI(4)P-coated Lipid Beads™ (Echelon Biosciences Inc.) pre-equilibrated with wash/binding buffer (10 mM HEPES, 0.25% NP-40, 150mM NaCl, pH 7.4) and incubated for 2 h at 4°C. The lysate was then removed, and the beads were washed with wash buffer three times for 5 min. The bound proteins were eluted by boiling in a 2×Laemmli buffer. Bound proteins were run, briefly, on SDS-PAGE 12% TGX stain-free gel. The gel was silver stained according to manufacturer instructions (Sigma), and the elution band was excised. The gel piece was then prepared for mass spectrometry analysis as previously described (82). Digested peptides were dried using a SpeedVac (ThermoFisher) and sent for mass spectrometry analysis using an Orbitrap Eclipse Tribid Mass Spectrometry (ThermoFisher). Subsequent results were analyzed using MaxQuant (Max Planck Institute of Biochemistry) with a false discovery rate of 1%.

### Protein-lipid overlay assay

Protein-lipid overlay assays were performed using commercially available PIP strip™ membranes (Echelon Biosciences Inc.). Nitrocellulose membranes prespotted with phospholipids were blocked with 2% nonfat milk in PBST (PBS and 0.1% Tween 20 (v/v), pH 7.5) for 1 h at room temperature. The blocked membranes were incubated with indicated purified GST-fusion protein (0.4-10 μg/mL in blocking buffer) for 1 h at room temperature. Protein binding to lipids was visualized with an anti-GST-HRP conjugated antibody (1:2,000) using a ChemiDoc Touch Imaging system (Bio-Rad).

### Subcellular localization assay and confocal microscopy

HeLa cells were transiently co-transfected for 16-20 h with mCherry and GFP-tagged constructs as listed in **SI Appendix, Table S2** using Lipofectamine 3000 transfection reagent (ThermoFisher Scientific). Cells were fixed in PBS with 4% paraformaldehyde for 20 min at room temperature, and coverslips were mounted using ProLong diamond anti-fade mountant (ThermoFisher Scientific). Images were acquired with a Zeiss LSM 880 laser-scanning confocal microscope using a 63× Plan-Apochromat objective lens (numerical aperture of 1.4) and operated with ZEN software (Carl Zeiss, Inc).

### Colocalization, quantification, and statistical analysis

For each condition, we acquired confocal images for 21 cells from 3 biological replicates. Quantitative colocalization analysis was performed using Imaris 10.2.0 (Oxford Instruments). mCherry and GFP puncta were segmented using machine learning in the Surfaces Creation Wizard. Eight representative images were used to train the machine learning algorithm. Masks of the segmented puncta were then generated, and colocalization was assessed by measuring their Mander’s overlap coefficients, which quantify the fraction of red pixels overlapping with green pixels. Shapiro-Wilk parametric hypothesis tests^65^ revealed that many of the data sets were non-normally distributed (**Supplementary Data S2**); therefore, statistical comparisons were made using Wilcoxon rank sum tests. Comparisons using the two-sample tests are also shown in **Supplementary Data S2** and similar results were obtained.

### Fold analysis with Flexible structure AlignmenT by Chaining Aligned fragment pairs allowing Twists (FATCAT)

3D models of effector proteins were retrieved from AlphaFold or global *Legionella* atlas database (https://godziklab.github.io/legionella-effectors/proteins). AlphaFold structures were cut to remove non-helical or non-PIP-binding regions of the protein. Helical regions or known PIP-binding *Legionella* domains were then used to perform a database search on FATCAT against: 1) the *Legionella* database of 660 *Legionella* domains and 2) the virulence factor database (vfdb) of 200 amino acid fragments of AlphaFold models of proteins from vfdb. Full results are shown in **Supplementary Data S3 & 4**.

### Structural similarity analysis and dendrogram construction

To assess structural relationships among PIP-binding effectors, we performed an all-against-all structural alignment using the DALI server ^66^. AlphaFold structures were submitted individually to DALI, and pairwise Z-scores were computed for all effector pairs. The resulting distance matrix was exported in Newick format, where higher structural similarity corresponds to shorter branch lengths. The Newick tree was imported into Geneious Prime (Biomatters Ltd.), where it was rendered and edited for clarity. Tree topology reflects relative structural similarity based on DALI Z-scores, and labels were manually curated for consistency with protein identifiers used throughout the manuscript.

## Acknowledgments

This work was supported by the NSF under CAREER Award IOS1750742, the NIH (NIAID, R21AI142317) to M.R.N and (NIAID/NIGMS, R01AI171196) to K.R.S., as well as the Delaware COBRE Program through NIGMS (P20GM104316, P30GM110758), which also supported access to the instrumentation at the Mass Spectrometry Facility. The content is solely the responsibility of the authors and does not necessarily represent the official views of the National Institutes of Health. We thank Jeff Caplan and the Delaware Biotechnology Institute Bioimaging Center, where the LSM880 confocal microscope at the University of Delaware was acquired with a shared instrumentation grant (S10 OD016361). Access to the instrumentation at the Bioimaging Center was partly supported by the DE-INBRE Core Facility Voucher Program with funding from the NIH (NIGMS, P20 GM103446) and the State of Delaware. M.R.N. would also like to thank the NIH for support through the Chemistry-Biology Interface (CBI) training grant, T32GM133395. Figure 1 was created using BioRender. We thank the Godzik lab for providing AlphaFold PDB files of the virulence factor database structures.

## Author Contributions

A.E.B. and M.R.N. conceived the study. A.E.B. performed all experiments with assistance from A.E.R., C.M.P., and R.R.N. for molecular cloning and protein-lipid overlay assays. Y.Y. performed mass spectrometry analysis. J.P.N. conducted statistical analysis. S.L.M. carried out colocalization analysis. K.R.S. performed bioinformatic and structural analyses. A.E.B., K.R.S., and M.R.N. analyzed and interpreted the data. M.R.N. supervised the study and secured funding. A.E.B. and M.R.N wrote the manuscript, and all authors provided editorial input.

