## Supplemental figures and tables for "Systematic Discovery of Novel Phosphoinositide-Binding Effectors in *Legionella* Reveals Conserved α-Helical Folds"

**This PDF file includes:**

Figures S1 to S7

Tables S1 to S3

SI References

**Other supporting materials for this manuscript include the following:**

Data S1: Pool maps and MS results

Data S2: Mander’s Overlap Coefficient analysis

Data S3: FATCAT results *Legionella* effector analysis

Data S4: FATCAT results *Coxiella* effector analysis

Supplementary Information: Figures

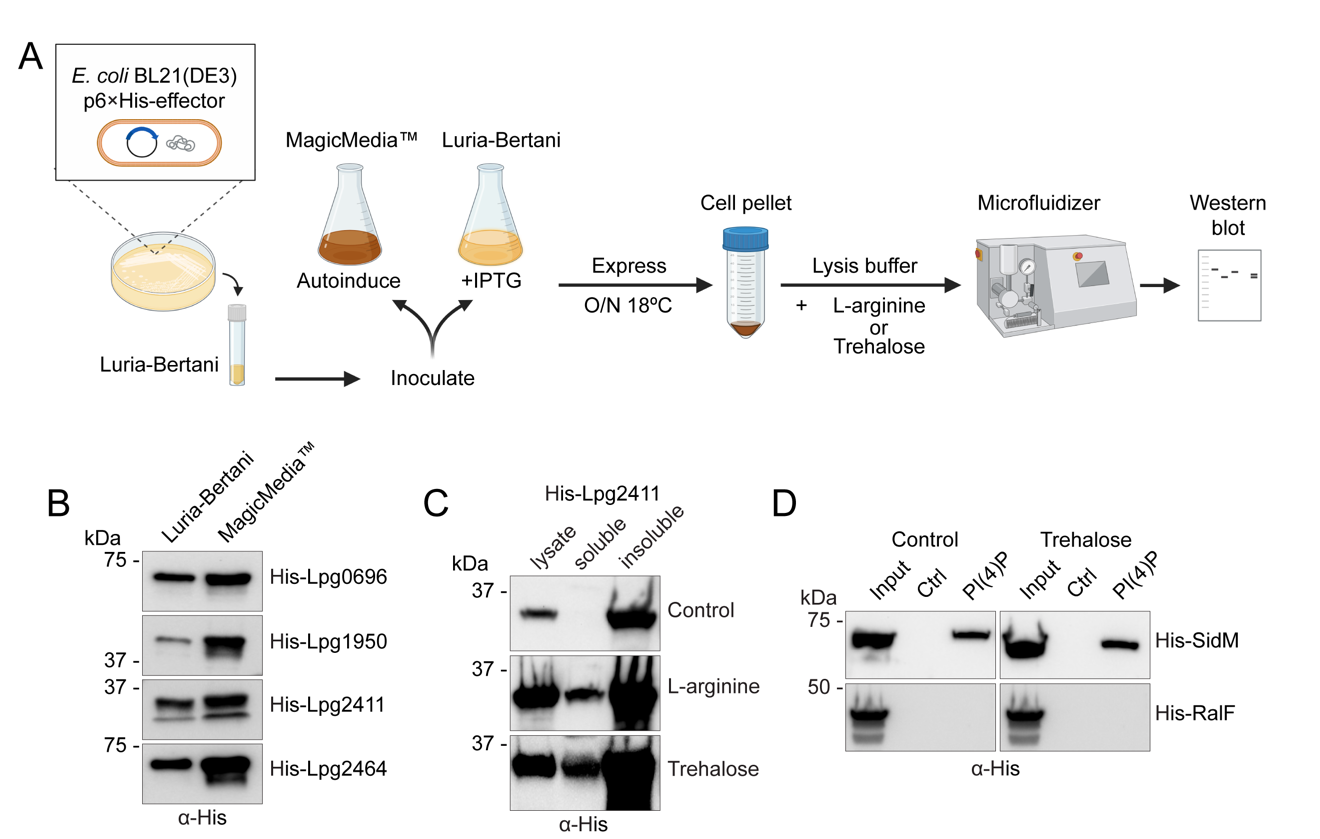

Fig. S1. Optimization of His-tagged effector protein expression and stability. (A) Workflow for effector expression and stability optimization. (B) Cultures of *E. coli* BL21(DE3) strains carrying plasmids encoding 6×His-tagged effectors were inoculated from overnight starter cultures in Luria-Bertani (LB) or MagicMedia™ and grown at 37°C. When an OD_600_ ~0.6 was reached, IPTG was added to the LB culture and both cultures were subsequently grown at 18°C overnight. Samples were normalized by volume, separated by SDS-PAGE, transferred to PVDF membrane, and probed with anti-His (mouse, 1:3,000). 6×His-Lpg0696/Lem3 ~65 kDa; 6×His-Lpg1950/RalF ~42 kDa; 6×His-Lpg2411/Lem24 ~33 kDa; 6×His-Lpg2464/SidM ~73 kDa (C) *E. coli* BL21(DE3) cells expressing 6×His-Lpg2411/Lem24 were collected after overnight growth at 18°C in LB. Cell pellets were resuspended in PBS-MM, PBS-MM + 300 mM L-arginine or PBS-MM + 750 mM trehalose and subsequently lysed with a microfluidizer. Whole cell lysate, soluble and insoluble fraction samples were separated by SDS-PAGE, transferred to PVDF membrane, and probed with anti-His (mouse, 1:3,000). (D) Proof of concept PIP-bead pulldown. *E. coli* BL21(DE3) lysate expressing 6×His-SidM or 6×His-RalF were incubated with agarose control beads or PI(4)P-coated beads. After incubation, beads were washed and elutions were separated via SDS-PAGE, transferred to PVDF membrane, and detected with anti-His (mouse, 1:3,000). 6×His-SidM ~ 73kDa; 6×His-RalF ~42 kDa.

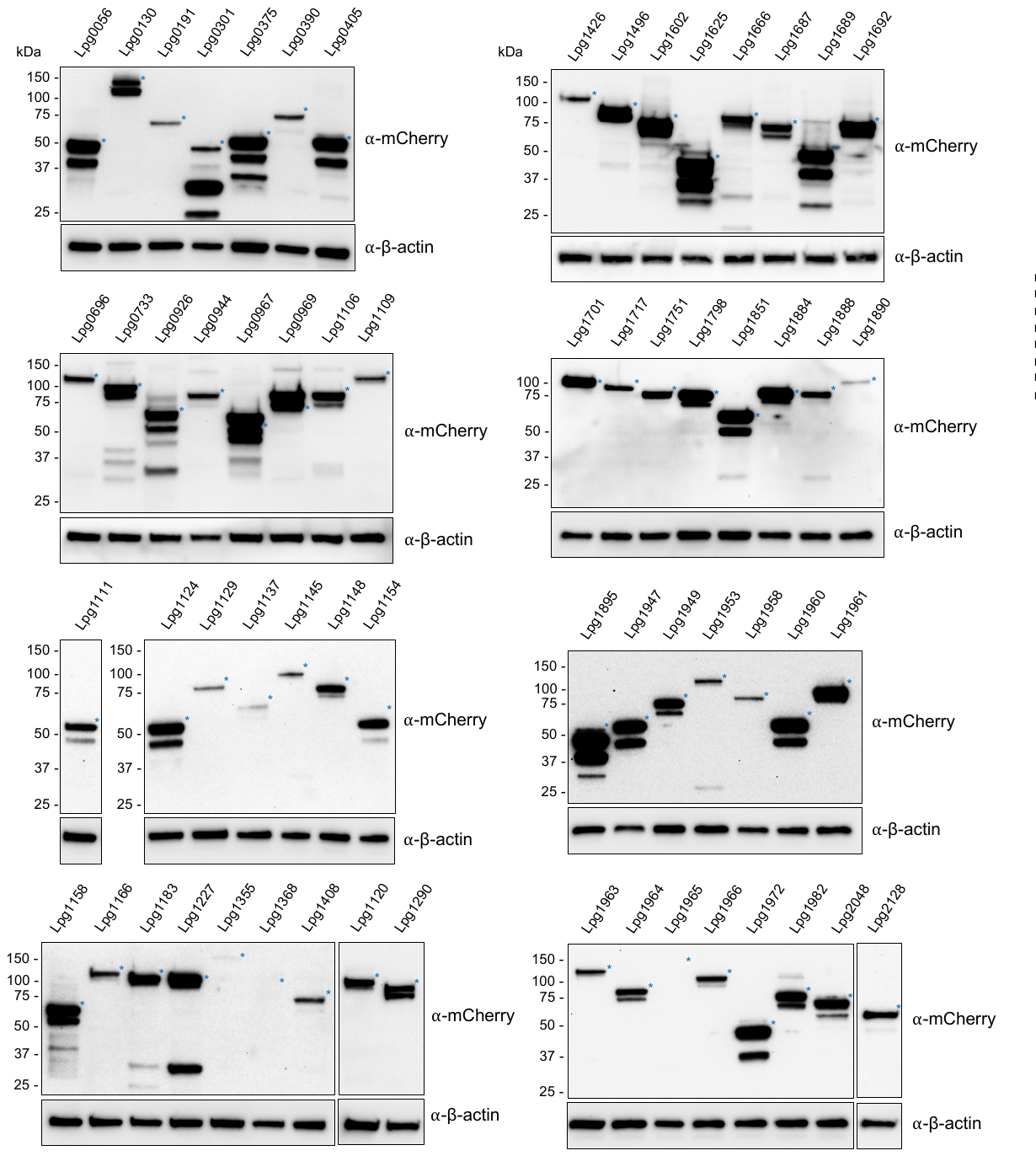

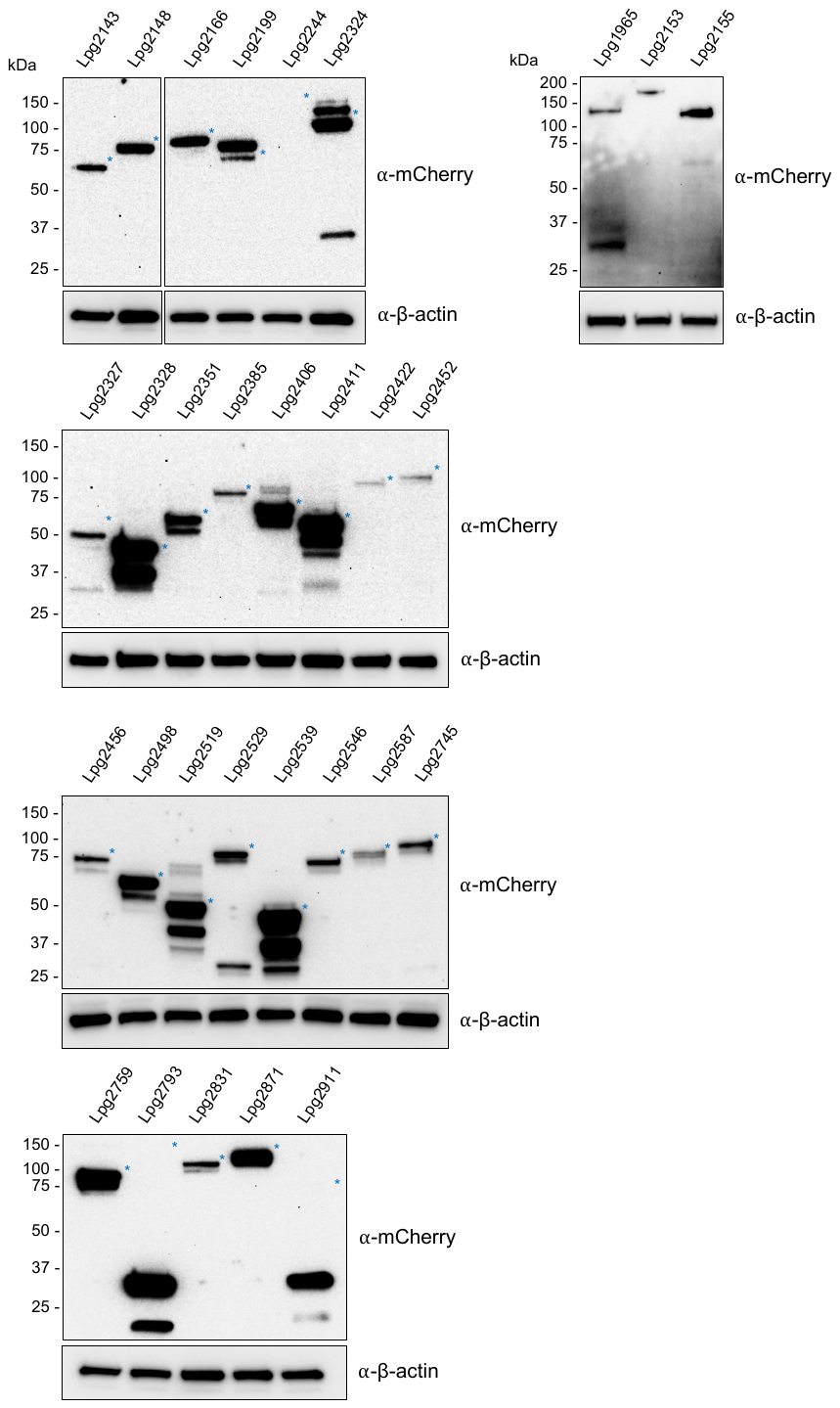
Fig. S2. mCherry-tagged effector fusion construct expression. HeLa cells transiently transfected with plasmids encoding mCherry-tagged effectors were lysed, lysate was separated via SDS-PAGE, transferred to a PVDF membrane, and probed with anti-mCherry (1:2,000) and anti-β-actin (1:2,000). Asterisks denote expected size of fusion construct.

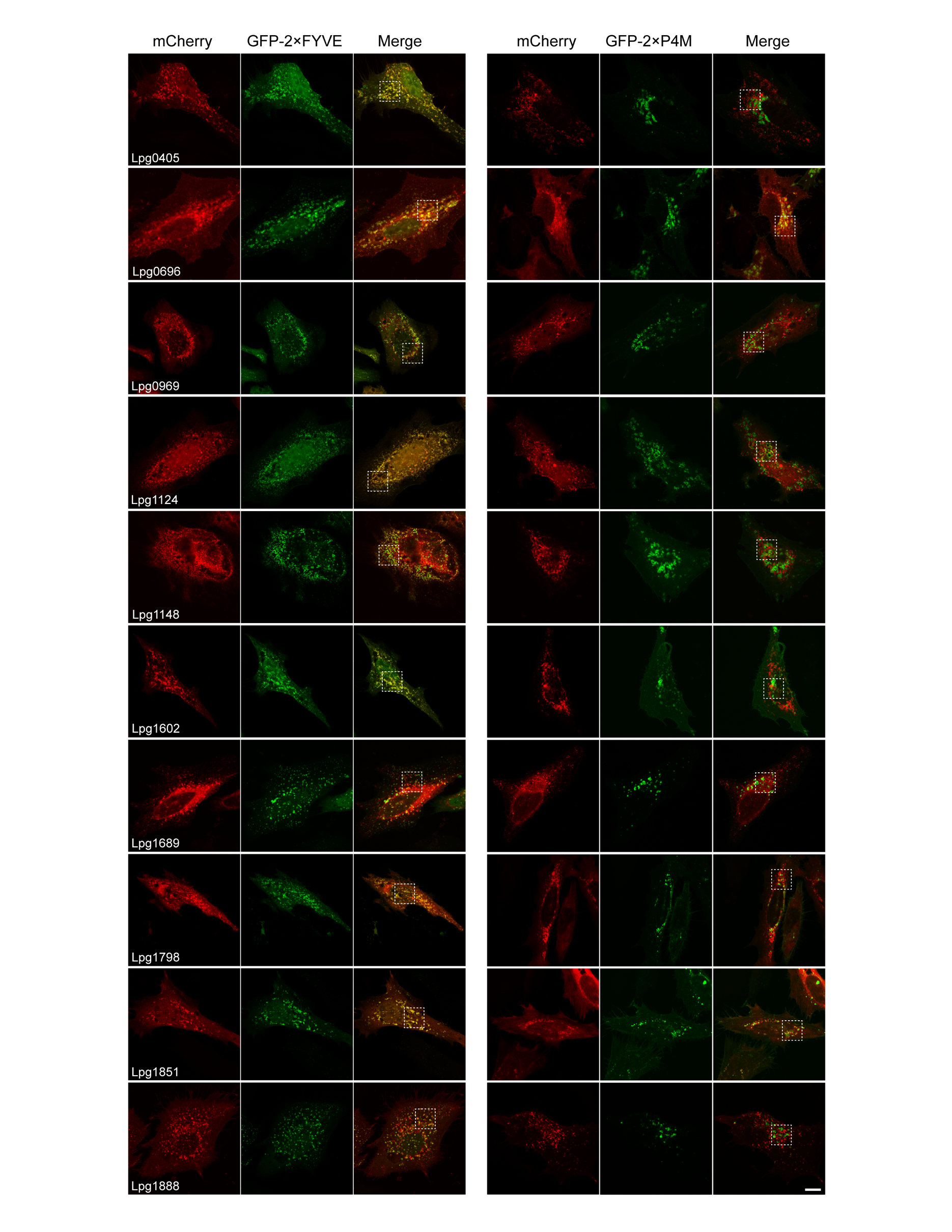

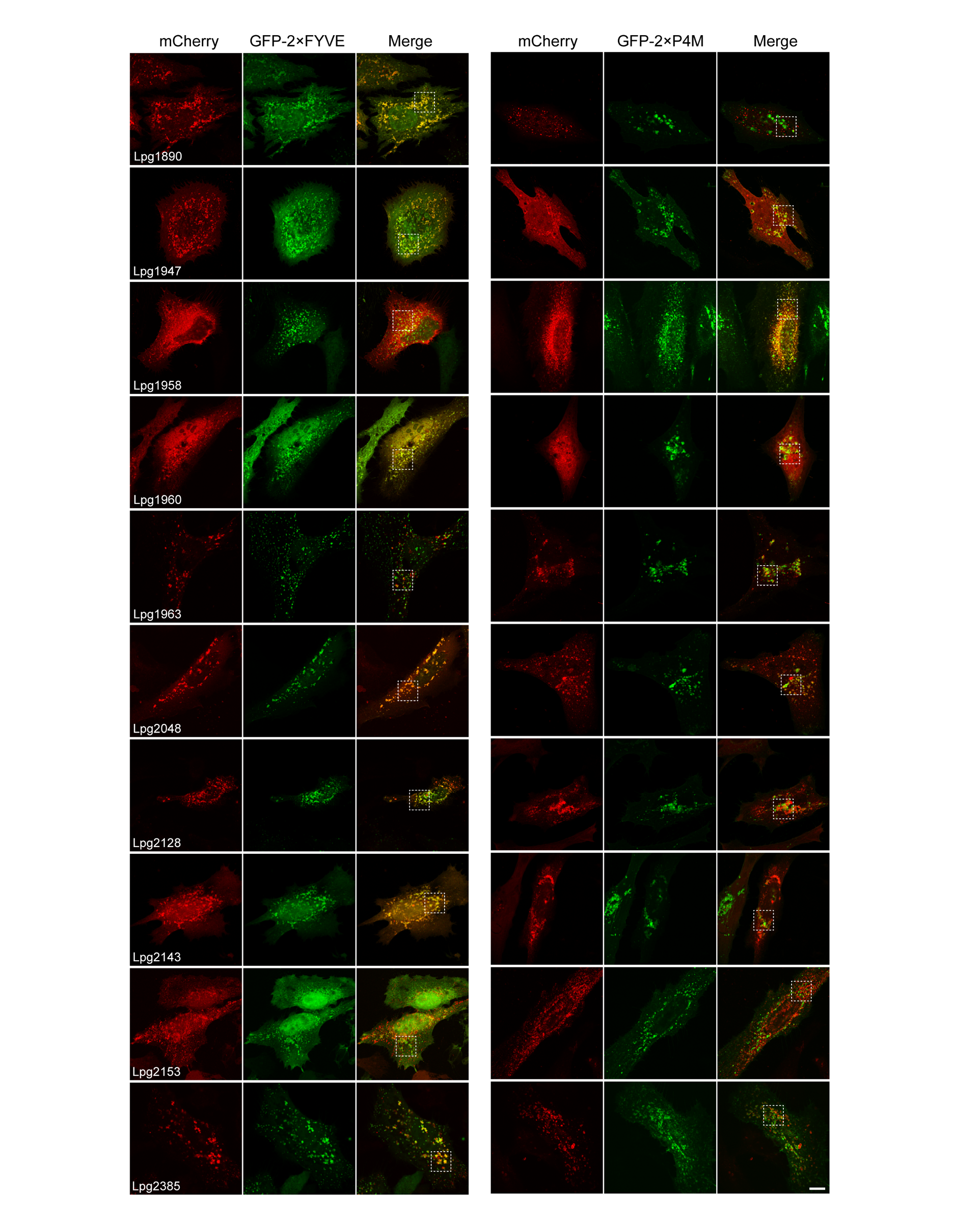

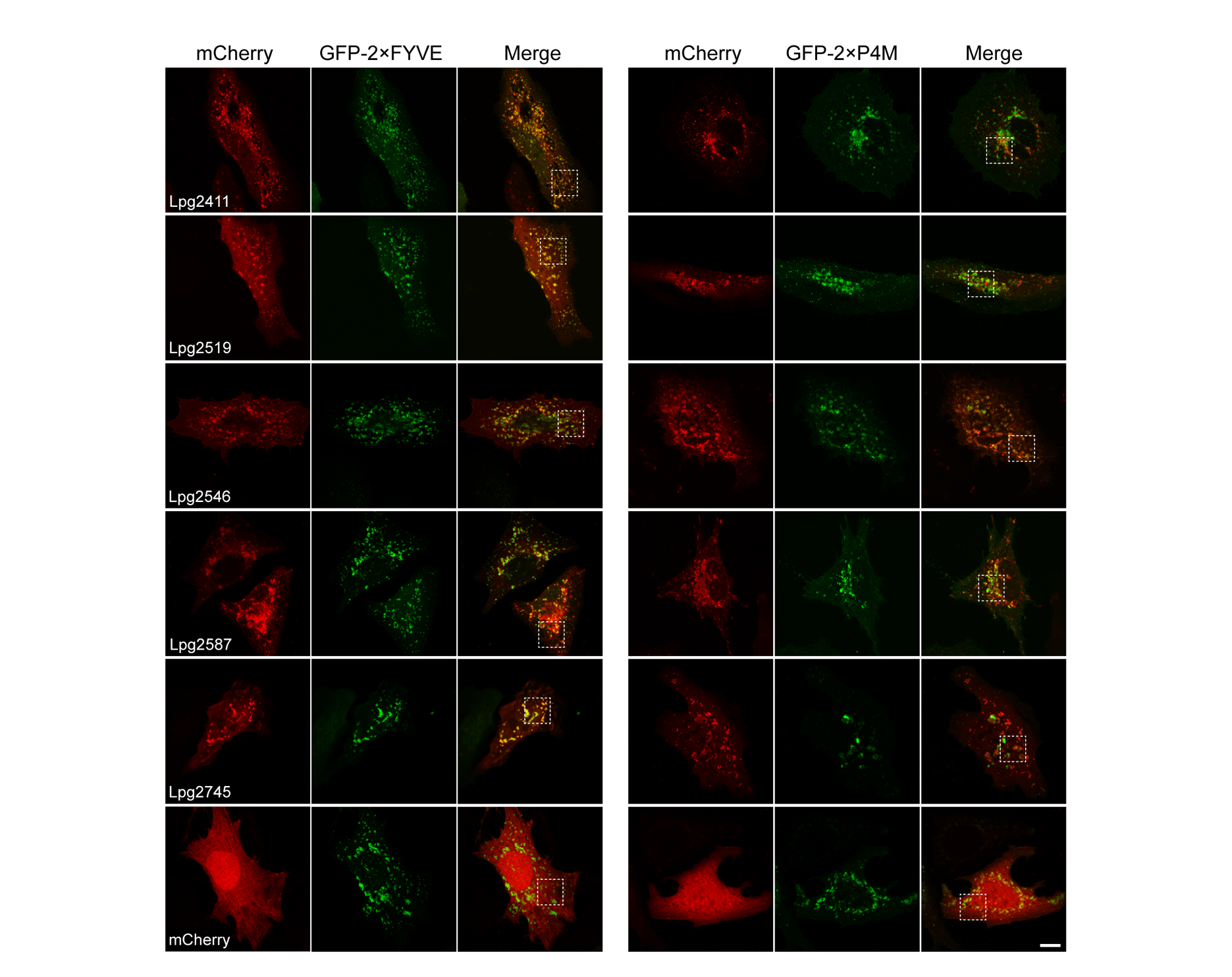

Fig. S3. Confocal images of mCherry-tagged effectors co-localizing with PIP-biosensor(s).

HeLa cells were transiently co-transfected with plasmids encoding mCherry-tagged effectors and GFP-tagged PIP-biosensors, GFP-2×FYVE or GFP-2×P4M, were visualized with confocal microscopy. Regions of interest shown in Figure 2 are indicated by white box. Scale bars, 5µm.

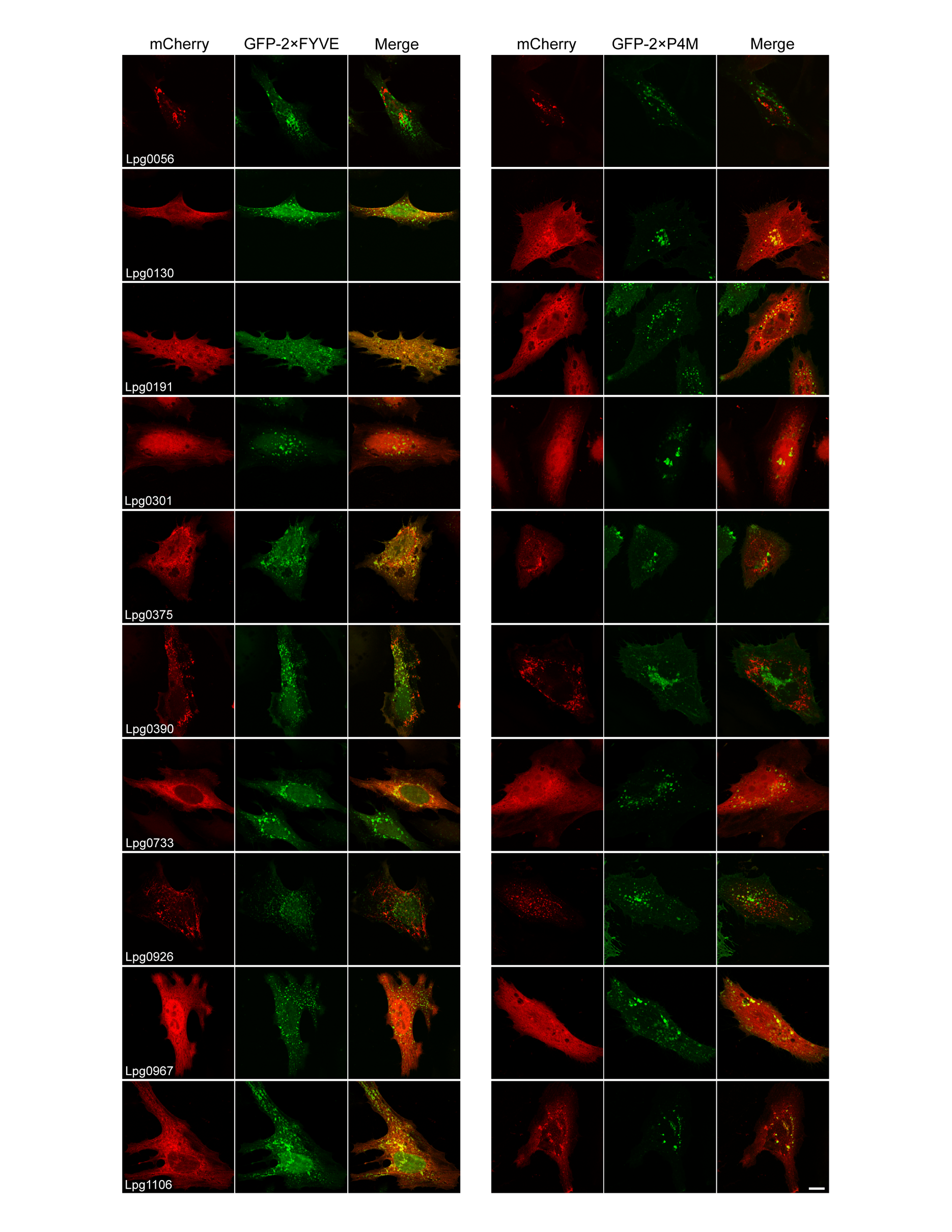

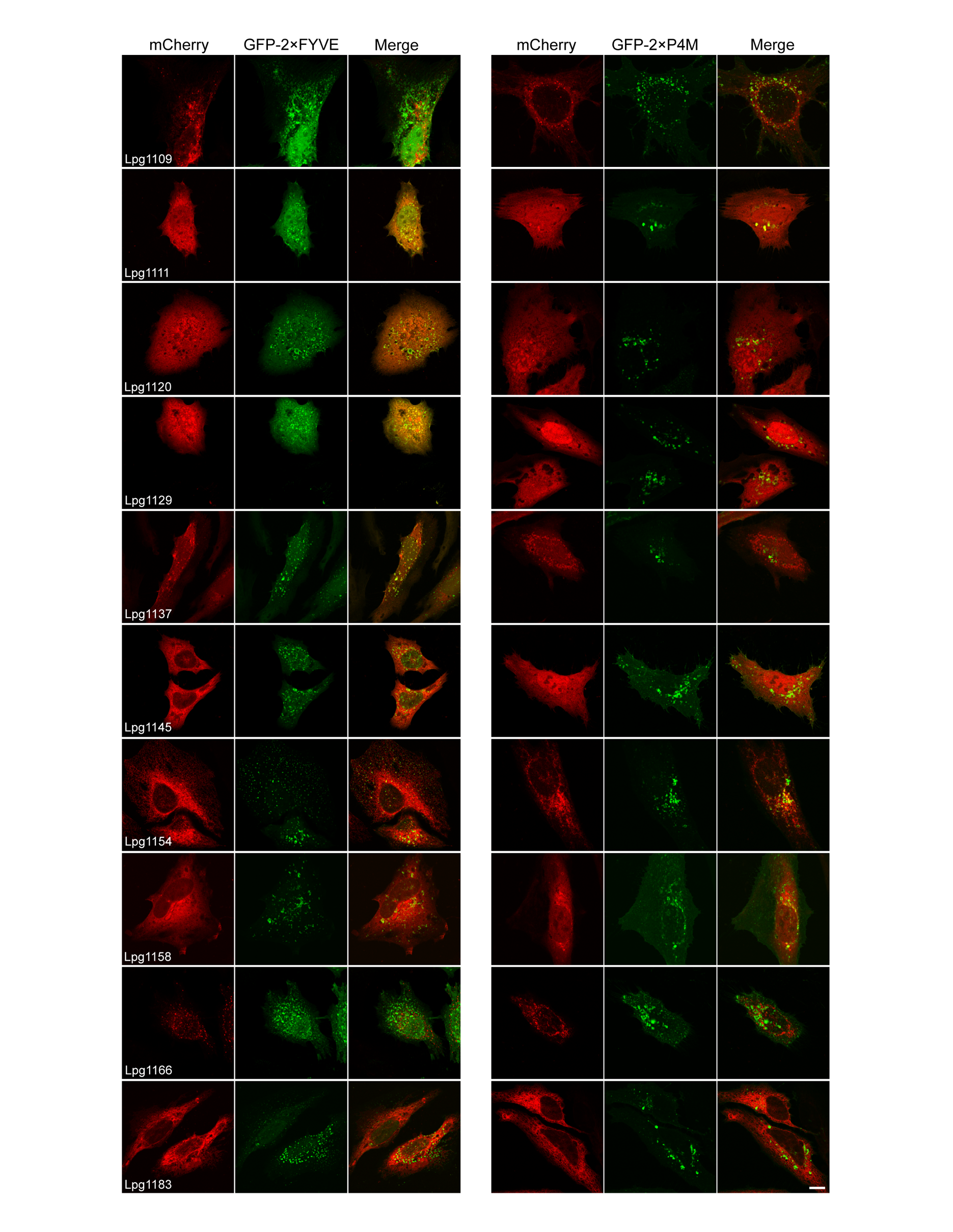

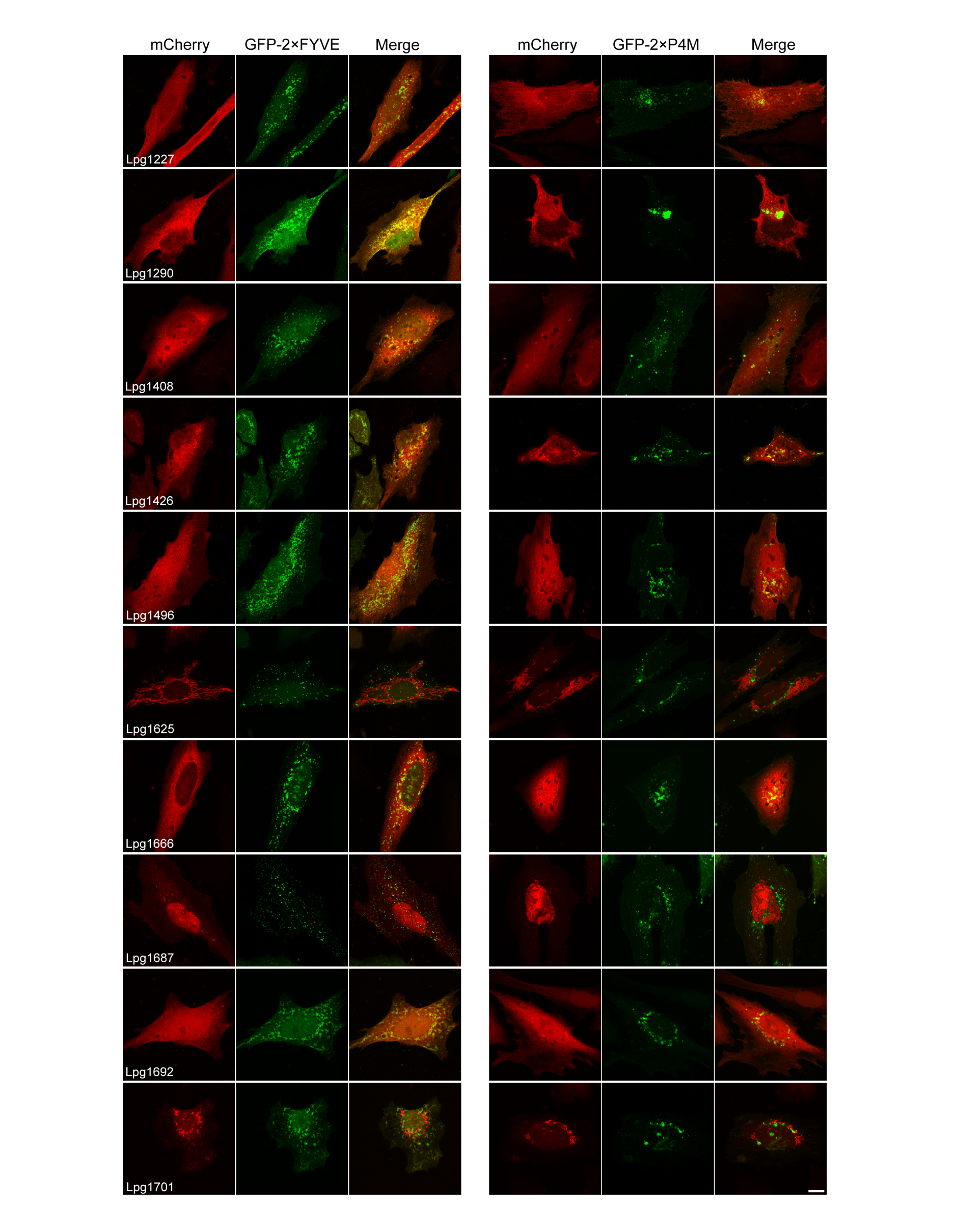

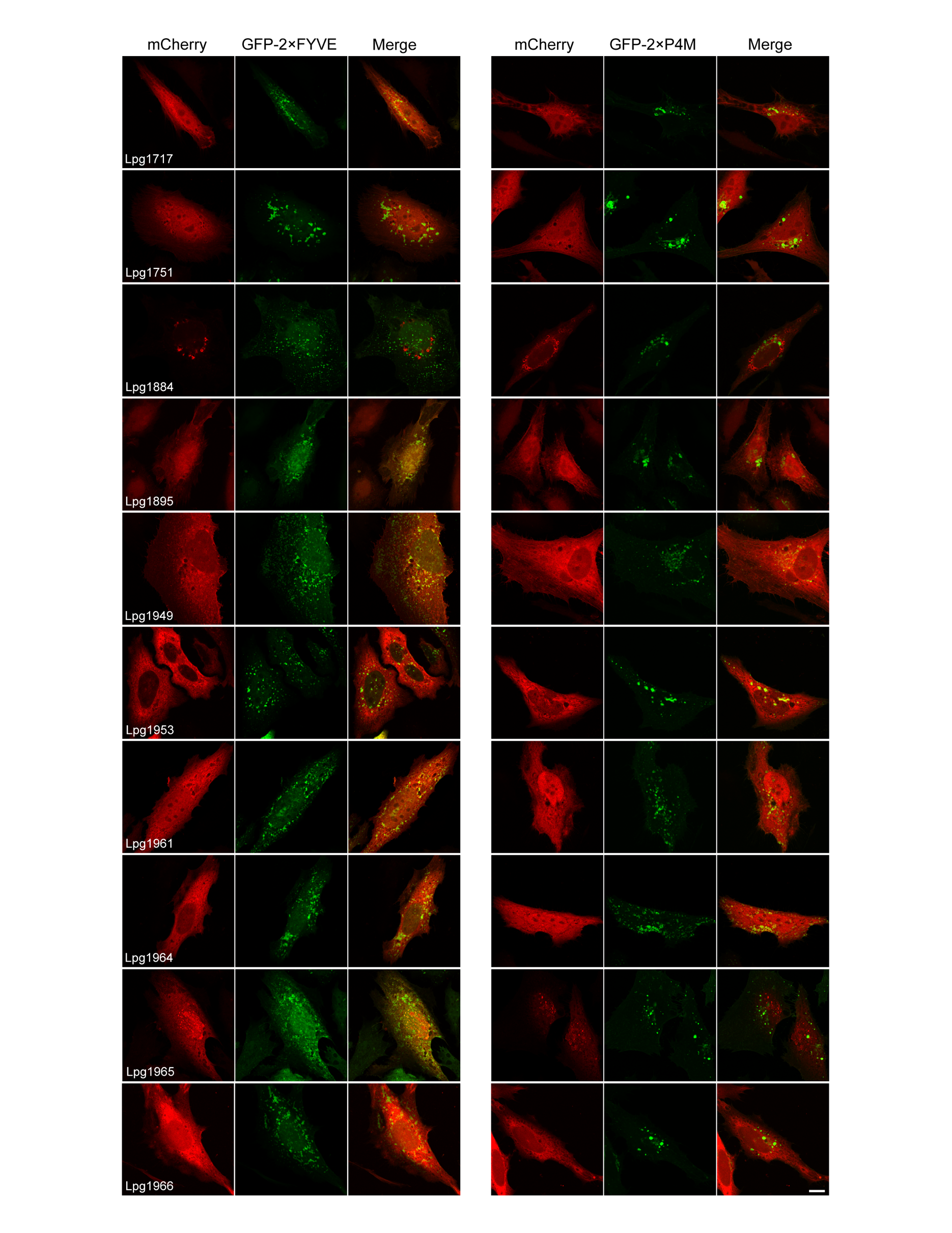

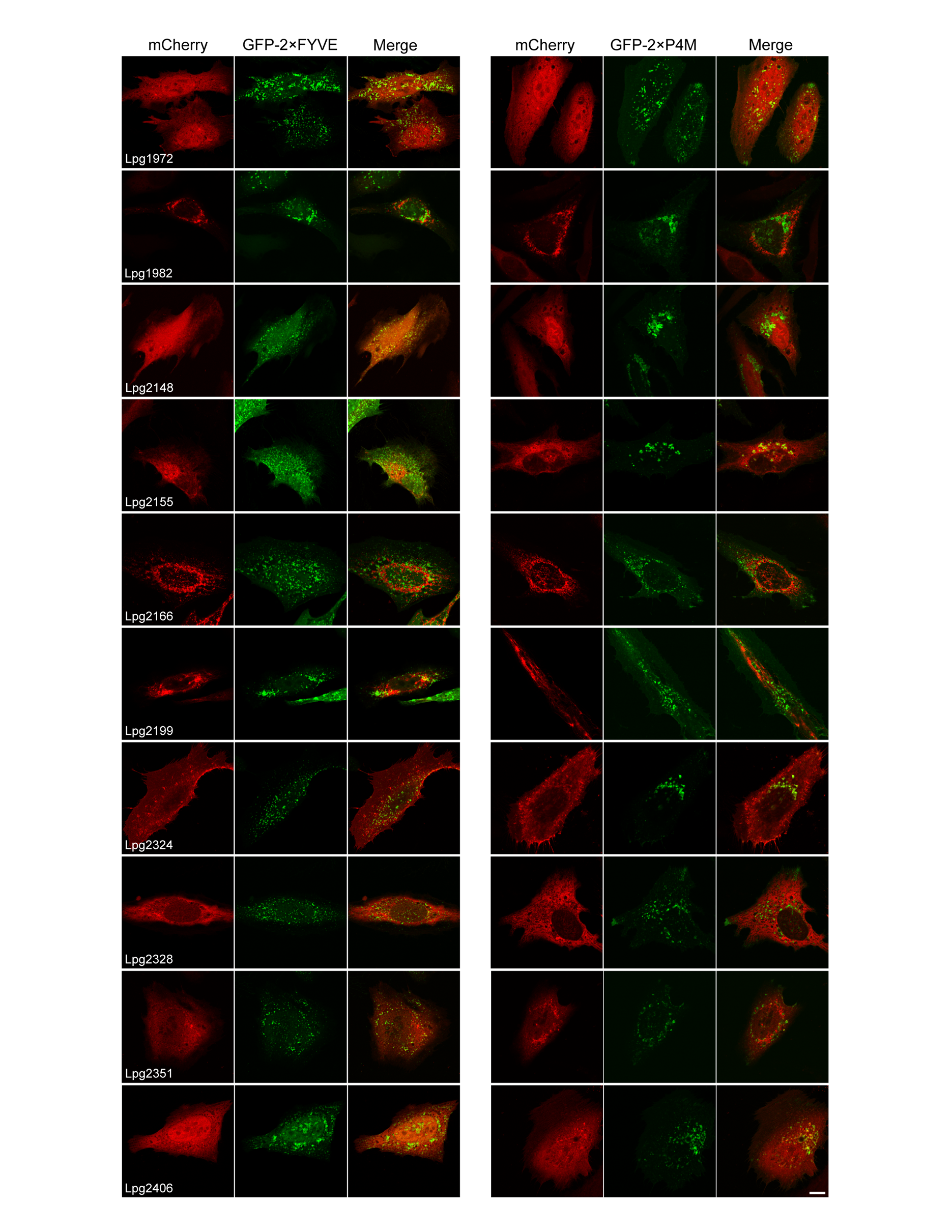

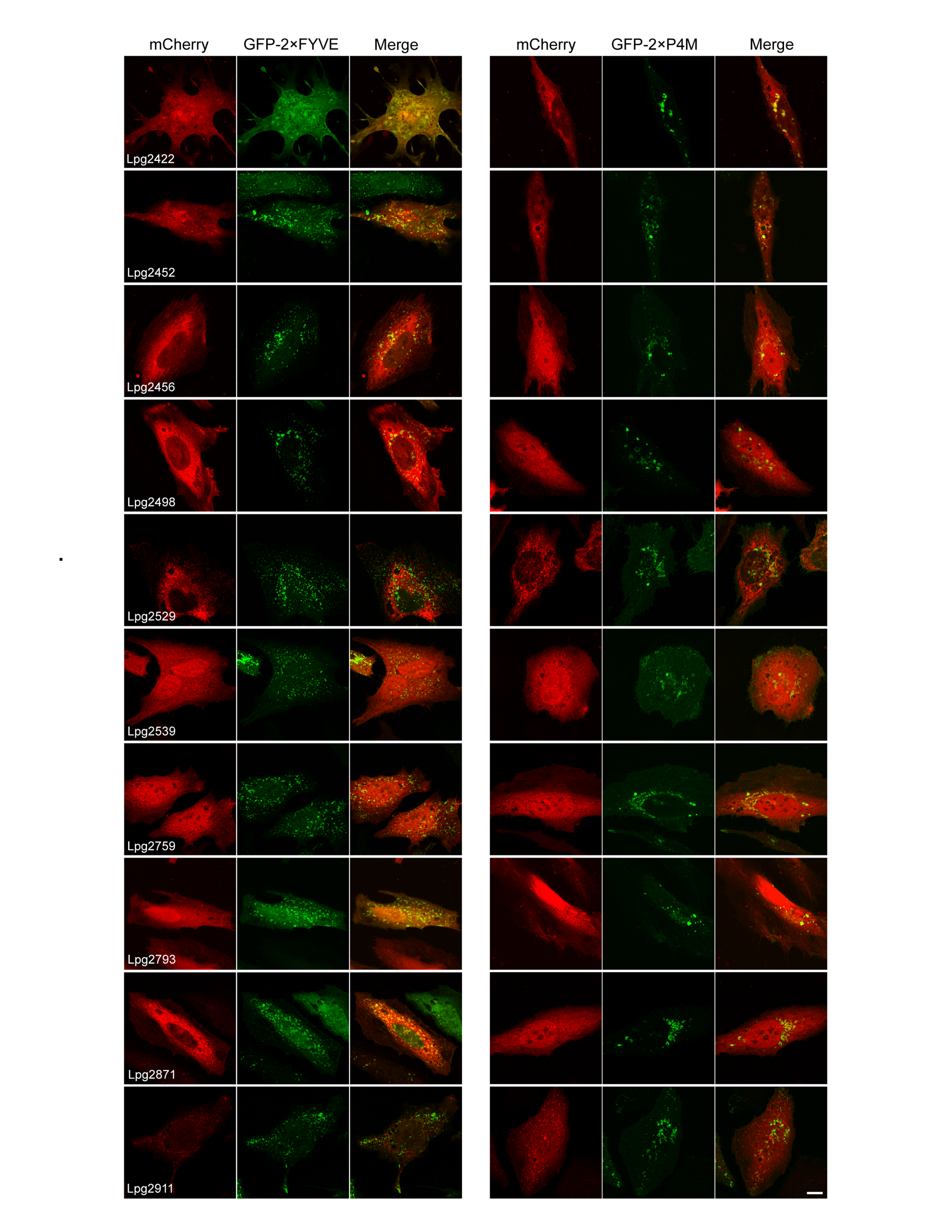

Fig. S4. Confocal images of mCherry-tagged effectors that do not localize with PIP-biosensors. HeLa cells were transiently co-transfected with plasmids encoding mCherry-tagged effectors and GFP-tagged PIP-biosensors, GFP-2×FYVE or GFP-2×P4M, were visualized with confocal microscopy. Scale bars, 5µm.

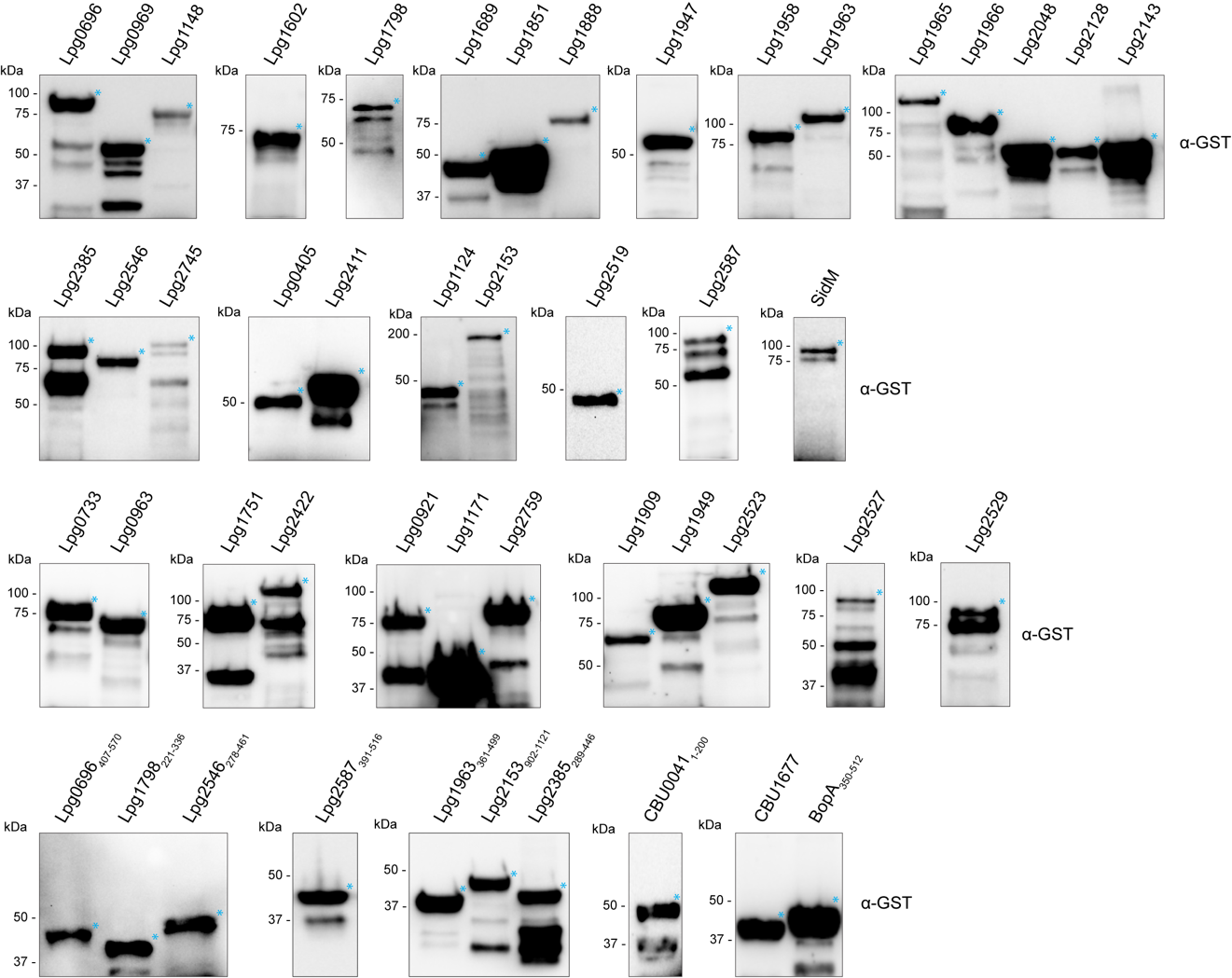

Fig. S5. Purified GST-fusion protein constructs. GST-fusions were expressed in *E. coli* BL21(DE3) strains, purified via affinity chromatography, separated via SDS-PAGE, transferred to a PVDF membrane, and probed with an anti-GST-HRP-conjugated antibody (1:3,000). Asterisks indicate the expected molecular weight of the full-length fusion proteins.

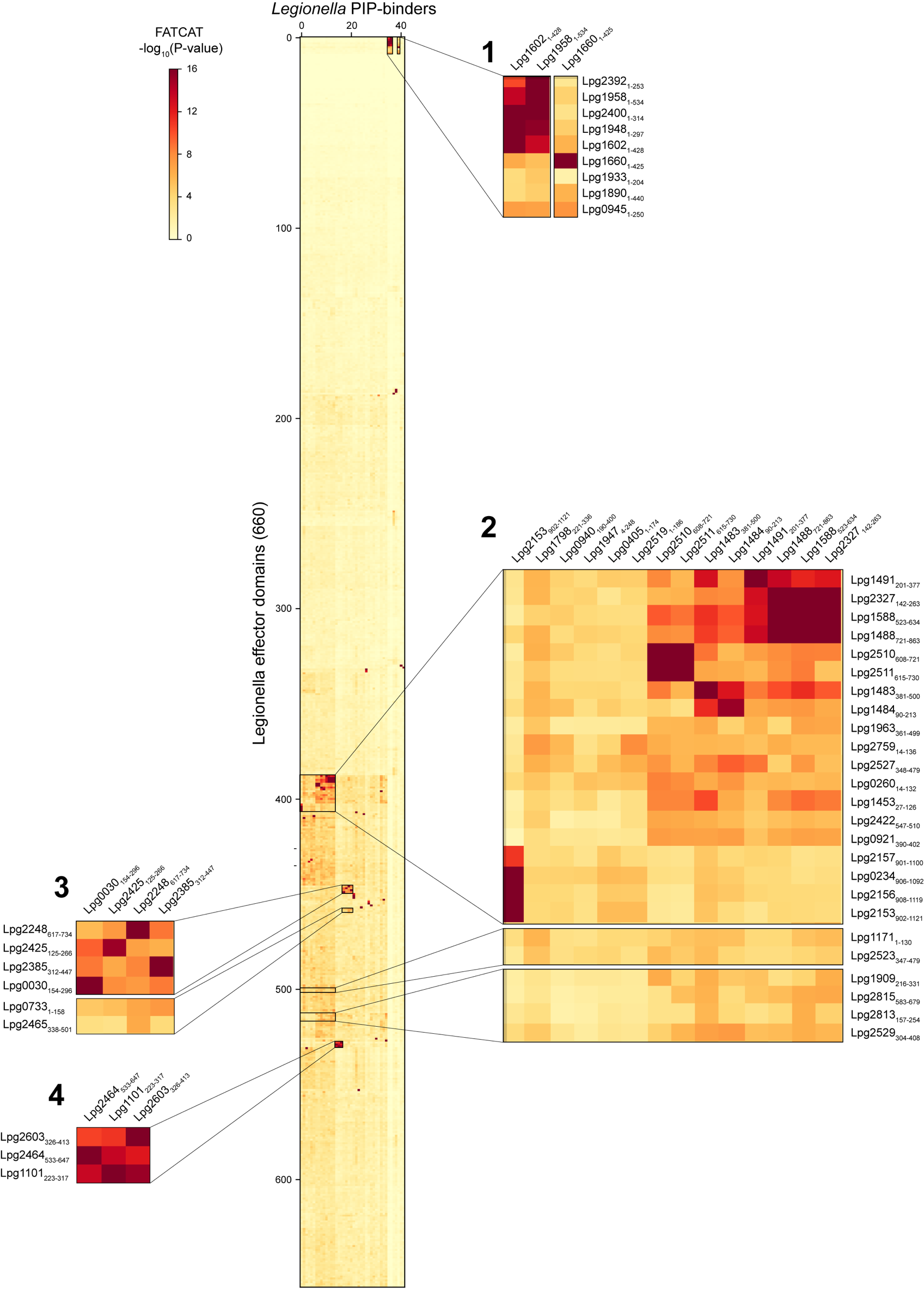

**Fig. S6. Asymmetric similarity matrix of the -log_10_(P-value) values from FATCAT analysis of *Legionella* PIP-binding effectors against *Legionella* effector domain database.** A matrix was generated using the -log_10_(P-value) from the comparison of PIP-binding effector domain structures (or full-length protein when the domain was unknown) to 660 *Legionella* effector domains by FATCAT. Axes are clustered by Ward’s method and selected areas of interest are magnified. Area 1 contains the Leucine-rich repeat containing effectors. Area 2 contains PIP-binding effectors with helical binding regions including P4C (Lpg2510/SidC, Lpg2511/SdcA) and LED006 domain-containing effectors (Lpg1483/LegK1, Lpg1484/Ceg22, Lpg1491/Lem9, Lpg1488/Lgt3, Lpg1588/LegC6, and Lpg2327) as well as additional PIP-binding effectors identified in this study and candidate PIP-binding effectors containing structurally similar helical folds. Area 3 contains the three effectors possessing LED035 (Lpg0030/RavB, Lpg2425/MavH, and Lpg2248/LotA) as well as the newly identified PIP-binding effector Lpg2385. Additionally, Lpg0733/RavH and Lpg2465/SidD cluster near this group. Area 4 contains the three effectors with the P4M domain (Lpg2464/SidM, Lpg1101, and Lpg2603).

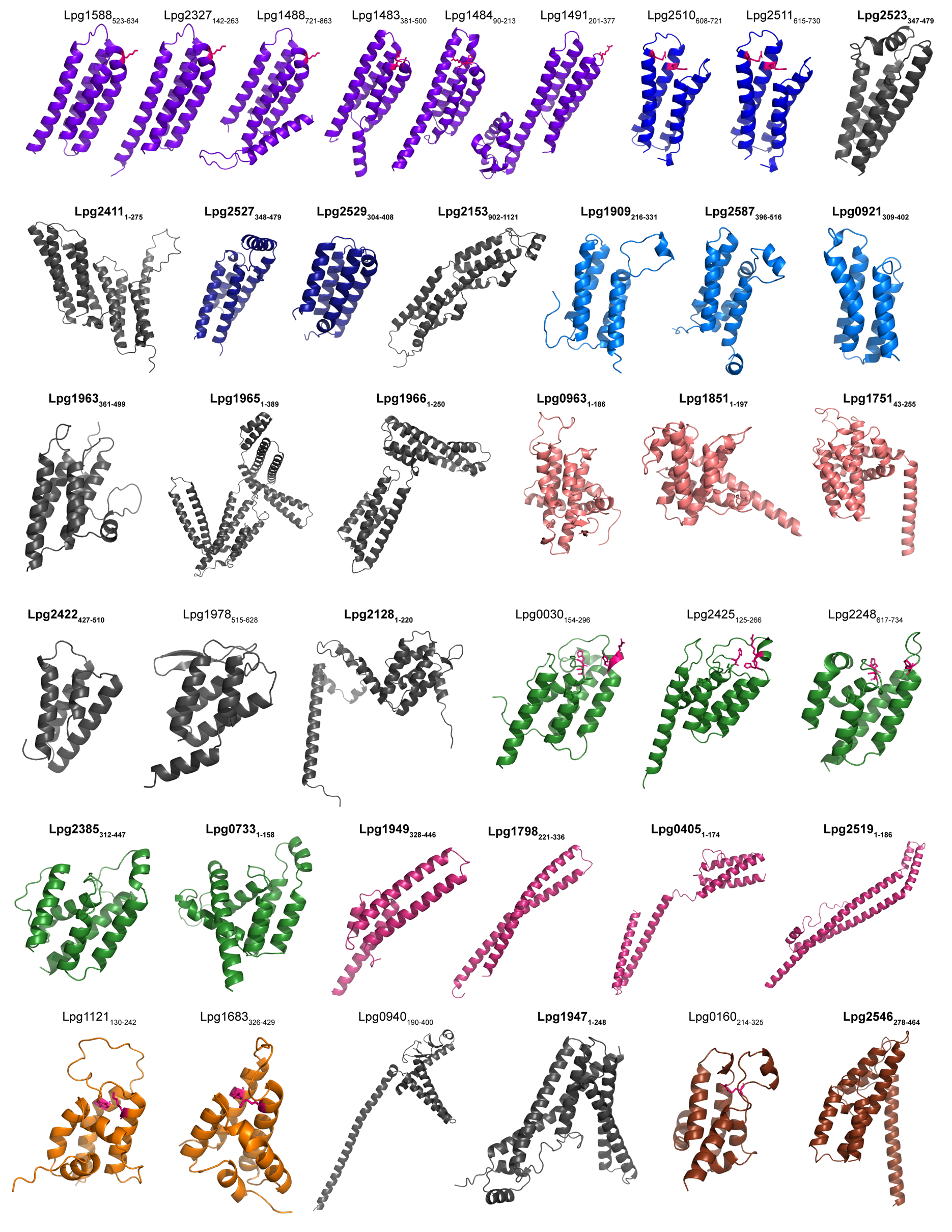

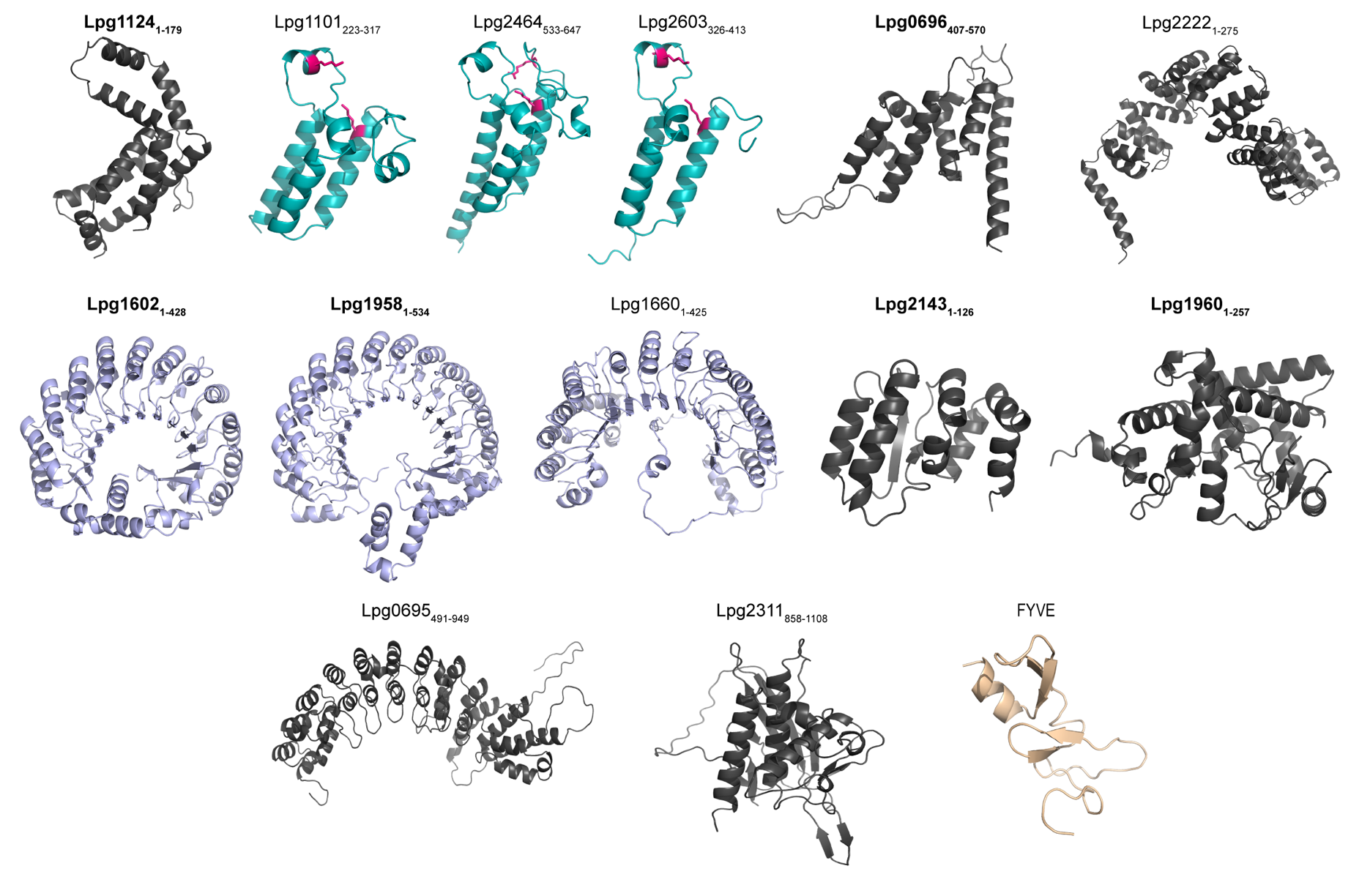

**Fig. S7. AlphaFold structures of PIP-binding effectors or effector domains included in the dendrogram analysis.** Structures are color-coded based on dendrogram groupings and branch color groupings in main figure 4. Labels in bold are PIP-binders discovered in this study. Known critical binding residues are depicted as pink sticks when applicable.

**Table S1. PIP-binding effector proteins**

| **PIP-binding effectors identified in this study:** | | | | | |
| --- | --- | --- | --- | --- | --- |
| **Effector Protein** | **PIP target** | **PIP-binding region** | **Fold description** | **Function** | **Ref.** |
| Lpg0405 | PI(3)P |  |  |  |  |
| Lpg0696  Lem3 | PI(3)P | 407-570 | 7 α-helix | Dephosphocholinates Rab1, removing AnkX modification | ^1^ |
| Lpg0733  RavH | PI(3)P | 1-158 | 5 α-helix bundle |  |  |
| Lpg0921  MavT | PI(3)P | 309-402 | 4 α-helix bundle |  |  |
| Lpg0963 | PI(3)P | 1-186 | Lpg1851 fold |  | ^2^ |
| Lpg1124 | PI(3)P |  |  |  |  |
| Lpg1602  LegL2 | PI(3)P |  | Leucine-rich repeat |  |  |
| Lpg1751 | PI(3)P | 43-255 | Lpg1851 fold |  | ^2^ |
| Lpg1798  MavU | PI(3)P | 221-336 | 2 α-helix anti-parallel |  |  |
| Lpg1851  Lem14 | PI(3)P | 1-197 | Small all α-helical protein | Synergistic with effector protein SidP, a PI3-phosphatase | ^3^ |
| Lpg1909  DenR | PI(3)P | 216-331 | 4 α-helix bundle | Recruits nRas to LCV resulting in reduced MAPK signaling | ^4^ |
| Lpg1947  Lem16 | PI(3)P |  |  |  |  |
| Lpg1949  Lem17 | PI(3)P | 328-446 | 3 α-helix | Possesses acetyltransferase activity | ^5^ |
| Lpg1958  LegL5 | PI(4)P  PI(5)P; PS |  | Leucine-rich repeat |  |  |
| Lpg1960  LirA | PI(3)P |  |  |  |  |
| Lpg1963  PieA | PI(3)P | 361-499 | 4 α-helix bundle | Localizes to LCV and disrupts lysosome morphology; on island of genomic plasticity | ^6^ |
| Lpg1965  PieC | PI(3)P | 1-389 |  | On island of genomic plasticity | ^6^ |
| Lpg1966  PieD | PI(3)P | 1-250 |  | On island of genomic plasticity | ^6^ |
| Lpg2128 | PI(3)P |  |  |  |  |
| Lpg2143 | PI(3)P |  |  |  |  |
| Lpg2153  SdeC | PI(3)P | 902-1121 | 10 α-helix bundle | Phosphoribosyl ubiquitination of host proteins including Rtn4 resulting in ER rearrangement | ^7-9^ |
| Lpg2385 | PI(3)P  PI(3,5)P_2_ | 312-447 | 5 α-helix bundle |  |  |
| Lpg2411  Lem24 | PI(3)P |  |  |  |  |
| Lpg2422  Lem25 | PI(3)P  PS |  | 4 α-helix bundle |  |  |
| Lpg2519 | PI(3)P  PI(5)P |  | 2 α-helix anti-parallel |  |  |
| Lpg2523  Lem26 | PI(3)P | 347-479 | 4 α-helix bundle |  |  |
| Lpg2527  LnaB | PI(3)P  PI(5)P | 348-479 | 4 α-helix bundle | Activates NF-κB; AMPylase of SidE generated PR-Ub, activity activated by actin binding | ^10-12^ |
| Lpg2529  Lem27 | PI(3)P | 304-408 | 4 α-helix bundle | Deubiquitinase; localizes to the LCV and regulated Rab10 ubiquitination levels | ^13^ |
| Lpg2546 | PI(3)P  PI(4)P | 278-464 | 7 α-helix bundle |  |  |
| Lpg2587 | PI(3)P | 391-516 | 4 α-helix bundle |  |  |
| **Previously known PIP-binding effectors:** | | | | | |
| **Effector Protein** | **PIP target** | **PIP-binding region** | **Fold description** | **Function** | **Ref.** |
| Lpg0030  RavB | PI(3)P | 721-863  LED035 | 5 α-helix bundle |  | ^14^ |
| Lpg0160  RavD | PI(3)P | 214-325 | 7 α-helix bundle | Deubiquitinase that removes ubiquitin from the LCV, prevents lysosomal fusion with LCV, and inhibits NF-κB signaling | ^15,16^ |
| Lpg0695  AnkX | PI(3)P; PI(4)P | 491-949 |  | Phosphocholinates Rab1 and 35; modulates endocytic traffic | ^17-20^ |
| Lpg0940  LidA | PI(3)P; PI(4)P | 190-400 |  | Facilitates Rab1 recruitment to the LCV and stabilizes active Rab1 | ^21-24^ |
| Lpg1101  Lem4 | PI(4)P | 223-317  P4M | 6 α-helix bundle | Phosphotyrosine phosphatase that localizes to lysosomes | ^25,26^ |
| Lpg1121  Ceg19 | PI(3)P | 13-242  LED027 | 5 α-helix bundle |  | ^14^ |
| Lpg1483  LegK1 | PI(3)P | 381-500  LED006 | 4 α-helix bundle | Activates NF-κB pathway through phosphorylation of IκB alpha | ^14,27^ |
| Lpg1484  Ceg22 | PI(3)P | 90-213  LED006 | 4 α-helix bundle |  | ^14^ |
| Lpg1488  LegC5/Lgt3 | PI(3)P | 721-863  LED006 | 4 α-helix bundle | Glycosyltransferase of eEF1A, inhibits protein translation | ^14,28,29^ |
| Lpg1491  Lem9 | PI(3)P | 201-377  LED006 | 4 α-helix bundle |  | ^14^ |
| Lpg1588  LegC6 | PI(3)P | 523-634  LED006 | 4 α-helix bundle |  | ^14^ |
| Lpg1660  LegL3 | PI(3)P; PS |  | Leucine-rich repeat |  | ^30^ |
| Lpg1683  RavZ | PI(3)P | 326-429  LED027 | 5 α-helix bundle | Inhibits autophagy through the irreversible deconjugation of Atg8 on autophagosomes | ^14,31,32^ |
| Lpg1978  SetA | PI(3)P | 615-728 | 4 α-helix bundle | Promiscuous glycosyltransferase activated by PI(3)P-binding; disrupts vesicular traffic | ^33-35^ |
| Lpg2222  LpnE | PI(3)P |  | ARM repeat | Interacts with OCLR1 on the LCV and assists with LCV biogenesis | ^36^ |
| Lpg2248  Lem21/LotA | PI(3)P | 617-734  LED035 | 5 α-helix bundle | Localizes to the LCV where it cleaves ubiquitin chains through deubiquitinase activity | ^14,37^ |
| Lpg2311  RidL | PI(3)P | 858-1108 |  | Binds to the retromer complex and inhibits retrograde traffic | ^38,39^ |
| Lpg2327 | PI(3)P | 348-479  LED006 | 4 α-helix bundle |  | ^14^ |
| Lpg2425  MavH | PI(3)P | 125-266  LED035 | 5 α-helix bundle | Interacts with actin capping protein to stimulate actin polymerization | ^14,40^ |
| Lpg2464  SidM | PI(4)P | 533-647  P4M | 6 α-helix bundle | Recruits and activates Rab1 through GEF activity at LCV, locks Rab1 in active state by AMPylation | ^21,24,41-47^ |
| Lpg2490  LepB | PI(3)P | 1210-1294  LED006 | 4 α-helix bundle | Rab1 GAP; PI4-kinase that generates PI(4)P on the LCV | ^14,47,48^ |
| Lpg2510  SdcA | PI(4)P | 608-721  P4C | 4 α-helix bundle | Paralog of Lpg2511 (SidC) | ^49-52^ |
| Lpg2511  SidC | PI(4)P | 615-730  P4C | 4 α-helix bundle | Recruits ER-derived vesicles to the LCV; E3 ubiquitin ligase | ^49-52^ |
| Lpg2603  Lem28 | PI(4)P | 326-413  P4M | 6 α-helix bundle | Kinase activated by inositol hexaphosphate | ^25,53^ |

**Table S2. Strains and plasmids used in this study**

| Strains or plasmid | Relevant features | Source or Reference |
| --- | --- | --- |
| ***E. coli* strains** |  |  |
| 2T1 | *F-mcrA Δ(mrr-hsdRMS-mcrBC) Φ80lacZΔM15 ΔlacX74 recA1 araΔ139 Δ(ara-leu)7697 galU galK rpsL (StrR) endA1 nupG fhuA::IS2* | ThermoFisher Scientific |
| GC5 | *F− ϕ80lacZΔM15 Δ(lacZYA-argF)U169 recA1 endA1 hsdR17(rK− mK+) phoA supE44 thi-1 gyrA96 relA1λ−tonA* | ThermoFisher Scientific |
| BL21 (DE3) | *F−ompT hsdS_B_(r_B_ − m_B_ − ) gal dcm(DE3)* | ThermoFisher Scientific |
| BL21 (DE3)pLysS | *F−ompT hsdS_B_(r_B_ − m_B_ − ) gal dcm(DE3) pLysS (CamR)* | ThermoFisher Scientific |
| **Plasmids** |  |  |
| ***E. coli* expression vectors** | | |
| pDONR221-effector library | Gateway entry vector with effectors; Kan^r^ | Machner, NIH |
| pDONR221-*lpg0696_407-560_* | pDONR221 carrying *lpg0696_407-560_* ; Kan^r^ | This study |
| pDONR221-*lpg1798_221-336_* | pDONR221 carrying *lpg1798_221-336_* ; Kan^r^ | This study |
| pDONR221-*lpg2153_903-1121_* | pDONR221 carrying *lpg2153_902-1121_ ;* Kan^r^ | This study |
| pDONR221-*lpg2385_289-447_* | pDONR221 carrying *lpg2385_289-447_ ;* Kan^r^ | This study |
| pDONR221-*lpg2546_278-461_* | pDONR221 carrying *lpg2546_278-464_ ;* Kan^r^ | This study |
| pDONR221-*lpg2587_391-516_* | pDONR221 carrying *lpg2587_391-516_ ;* Kan^r^ | This study |
| pDEST17-effector library | Encodes 6×His- effectors; Amp^r^ | Machner, NIH |
| pDEST15 | Encodes GST; Amp^r^ | ThermoFisher Scientific |
| pDEST15-*lpg0405* | Encoding IPTG-inducible expression of Lpg0405 with N-terminal GST tag; Amp^r^ | This study |
| pDEST15-*lpg0696* | Encoding IPTG-inducible expression of Lpg0696 with N-terminal GST tag; Amp^r^ | This study |
| pDEST15-*lpg0696_407-560_* | Encoding IPTG-inducible expression of Lpg0696_407-560_ with N-terminal GST tag; Amp^r^ | This study |
| pDEST15-*lpg0733* | Encoding IPTG-inducible expression of Lpg0733 with N-terminal GST tag; Amp^r^ | This study |
| pDEST15-*lpg0921* | Encoding IPTG-inducible expression of Lpg0921 with N-terminal GST tag; Amp^r^ | This study |
| pDEST15-*lpg0963* | Encoding IPTG-inducible expression of Lpg0963 with N-terminal GST tag; Amp^r^ | This study |
| pDEST15-*lpg0969* | Encoding IPTG-inducible expression of Lpg0969 with N-terminal GST tag; Amp^r^ | This study |
| pDEST15-*lpg1124* | Encoding IPTG-inducible expression of Lpg1124 with N-terminal GST tag; Amp^r^ | This study |
| pDEST15-*lpg1148* | Encoding IPTG-inducible expression of Lpg1148 with N-terminal GST tag; Amp^r^ | This study |
| pDEST15-*lpg1602* | Encoding IPTG-inducible expression of Lpg1602 with N-terminal GST tag; Amp^r^ | This study |
| pDEST15-*lpg1689* | Encoding IPTG-inducible expression of Lpg1689 with N-terminal GST tag; Amp^r^ | This study |
| pDEST15-*lpg1751* | Encoding IPTG-inducible expression of Lpg1751 with N-terminal GST tag; Amp^r^ | This study |
| pDEST15-*lpg1798* | Encoding IPTG-inducible expression of Lpg1798 with N-terminal GST tag; Amp^r^ | This study |
| pDEST15-*lpg1798_221-336_* | Encoding IPTG-inducible expression of Lpg1798_221-336_ with N-terminal GST tag; Amp^r^ | This study |
| pDEST15-*lpg1851* | Encoding IPTG-inducible expression of Lpg1851 with N-terminal GST tag; Amp^r^ | This study |
| pDEST15-*lpg1888* | Encoding IPTG-inducible expression of Lpg1888 with N-terminal GST tag; Amp^r^ | This study |
| pDEST15-*lpg1890* | Encoding IPTG-inducible expression of Lpg1890 with N-terminal GST tag; Amp^r^ | This study |
| pDEST15-*lpg1909* | Encoding IPTG-inducible expression of Lpg1909 with N-terminal GST tag; Amp^r^ | This study |
| pDEST15-*lpg1947* | Encoding IPTG-inducible expression of Lpg1947 with N-terminal GST tag; Amp^r^ | This study |
| pDEST15-*lpg1949* | Encoding IPTG-inducible expression of Lpg1949 with N-terminal GST tag; Amp^r^ | This study |
| pDEST15-*lpg1958* | Encoding IPTG-inducible expression of Lpg1958 with N-terminal GST tag; Amp^r^ | This study |
| pDEST15-*lpg1960* | Encoding IPTG-inducible expression of Lpg1960 with N-terminal GST tag; Amp^r^ | This study |
| pDEST15-*lpg1963* | Encoding IPTG-inducible expression of Lpg1963 with N-terminal GST tag; Amp^r^ | This study |
| pDEST15-*lpg1965* | Encoding IPTG-inducible expression of Lpg1965 with N-terminal GST tag; Amp^r^ | This study |
| pDEST15-*lpg1966* | Encoding IPTG-inducible expression of Lpg1966 with N-terminal GST tag; Amp^r^ | This study |
| pDEST15-*lpg2048* | Encoding IPTG-inducible expression of Lpg2048 with N-terminal GST tag; Amp^r^ | This study |
| pDEST15-*lpg2128* | Encoding IPTG-inducible expression of Lpg2128 with N-terminal GST tag; Amp^r^ | This study |
| pDEST15-*lpg2143* | Encoding IPTG-inducible expression of Lpg2143 with N-terminal GST tag; Amp^r^ | This study |
| pDEST15-*lpg2153* | Encoding IPTG-inducible expression of Lpg2153 with N-terminal GST tag; Amp^r^ | This study |
| pDEST15-*lpg2153_902-1121_* | Encoding IPTG-inducible expression of Lpg2153_902-1121_ with N-terminal GST tag; Amp^r^ | This study |
| pDEST15-*lpg2385* | Encoding IPTG-inducible expression of Lpg2385 with N-terminal GST tag; Amp^r^ | This study |
| pDEST15-*lpg2385_289-447_* | Encoding IPTG-inducible expression of Lpg2385_289-447_ with N-terminal GST tag; Amp^r^ | This study |
| pDEST15-*lpg2411* | Encoding IPTG-inducible expression of Lpg2411 with N-terminal GST tag; Amp^r^ | This study |
| pDEST15-*lpg2422* | Encoding IPTG-inducible expression of Lpg2422 with N-terminal GST tag; Amp^r^ | This study |
| pDEST15-*lpg2519* | Encoding IPTG-inducible expression of Lpg2519 with N-terminal GST tag; Amp^r^ | This study |
| pDEST15-*lpg2523* | Encoding IPTG-inducible expression of Lpg2523 with N-terminal GST tag; Amp^r^ | This study |
| pDEST15-*lpg2527* | Encoding IPTG-inducible expression of Lpg2527 with N-terminal GST tag; Amp^r^ | This study |
| pDEST15-*lpg2529* | Encoding IPTG-inducible expression of Lpg2529 with N-terminal GST tag; Amp^r^ | This study |
| pDEST15-*lpg2546* | Encoding IPTG-inducible expression of Lpg2546 with N-terminal GST tag; Amp^r^ | This study |
| pDEST15-*lpg2546_278-464_* | Encoding IPTG-inducible expression of Lpg2546_278-464_ with N-terminal GST tag; Amp^r^ | This study |
| pDEST15-*lpg2587* | Encoding IPTG-inducible expression of Lpg2587 with N-terminal GST tag; Amp^r^ | This study |
| pDEST15-*lpg2587_391-516_* | Encoding IPTG-inducible expression of Lpg2587_391-517_ with N-terminal GST tag; Amp^r^ | This study |
| pDEST15-*lpg2745* | Encoding IPTG-inducible expression of Lpg2745 with N-terminal GST tag; Amp^r^ | This study |
| pGEX-6P-1-*lpg1963_361-499_* | Encoding IPTG-inducible expression of Lpg1963_361-499_ with N-terminal cleavable GST tag; Amp^r^ | GenScript |
| pGEX-6P-1-*cbu1677* | Encoding IPTG-inducible expression of CBU1677 with N-terminal cleavable GST tag; Amp^r^ | GenScript |
| pGEX-6P-1-*cbu0041_1-200_* | Encoding IPTG-inducible expression of CBU0041_1-200_ with N-terminal cleavable GST tag; Amp^r^ | GenScript |
| pGEX-6P-1-b*opA_350-512_* | Encoding IPTG-inducible expression of BopA_350-512_ with N-terminal cleavable GST tag; Amp^r^ | GenScript |
| **Mammalian expression vectors** | | |
| pEGFP-2×FYVE | Encoding GFP-tagged HRS tandem FYVE domain | ^54^ |
| EGFP-2×P4M | Encoding GFP-tagged tandem P4M domain, Kan^r^ | ^43^ |
| 362 pCS mCherry-DEST | Encodes mCherry; Cam^r^, Amp^r^ | ^55^ |
| 362 pCS Cherry-*lpg0056* | Encoding mCherry-tagged Lpg0056, Cam^r^, Amp^r^ | This study |
| 362 pCS Cherry-*lpg0130* | Encoding mCherry-tagged Lpg0130, Cam^r^, Amp^r^ | This study |
| 362 pCS Cherry-*lpg0191* | Encoding mCherry-tagged Lpg0191, Cam^r^, Amp^r^ | This study |
| 362 pCS Cherry-*lpg0301* | Encoding mCherry-tagged Lpg0301, Cam^r^, Amp^r^ | This study |
| 362 pCS Cherry-*lpg0375* | Encoding mCherry-tagged Lpg0375, Cam^r^, Amp^r^ | This study |
| 362 pCS Cherry-*lpg0390* | Encoding mCherry-tagged Lpg0390, Cam^r^, Amp^r^ | This study |
| 362 pCS Cherry-*lpg0405* | Encoding mCherry-tagged Lpg0405, Cam^r^, Amp^r^ | This study |
| 362 pCS Cherry-*lpg0696* | Encoding mCherry-tagged Lpg0696, Cam^r^, Amp^r^ | This study |
| 362 pCS Cherry-*lpg0733* | Encoding mCherry-tagged Lpg0733, Cam^r^, Amp^r^ | This study |
| 362 pCS Cherry-*lpg0926* | Encoding mCherry-tagged Lpg0926, Cam^r^, Amp^r^ | This study |
| 362 pCS Cherry-*lpg0944* | Encoding mCherry-tagged Lpg0944, Cam^r^, Amp^r^ | This study |
| 362 pCS Cherry-*lpg0967* | Encoding mCherry-tagged Lpg0967, Cam^r^, Amp^r^ | This study |
| 362 pCS Cherry-*lpg0969* | Encoding mCherry-tagged Lpg0969, Cam^r^, Amp^r^ | This study |
| 362 pCS Cherry-*lpg1106* | Encoding mCherry-tagged Lpg1106, Cam^r^, Amp^r^ | This study |
| 362 pCS Cherry-*lpg1109* | Encoding mCherry-tagged Lpg1109, Cam^r^, Amp^r^ | This study |
| 362 pCS Cherry-*lpg1111* | Encoding mCherry-tagged Lpg1111, Cam^r^, Amp^r^ | This study |
| 362 pCS Cherry-*lpg1120* | Encoding mCherry-tagged Lpg1120, Cam^r^, Amp^r^ | This study |
| 362 pCS Cherry-*lpg1124* | Encoding mCherry-tagged Lpg1124, Cam^r^, Amp^r^ | This study |
| 362 pCS Cherry-*lpg1129* | Encoding mCherry-tagged Lpg1129, Cam^r^, Amp^r^ | This study |
| 362 pCS Cherry-*lpg1137* | Encoding mCherry-tagged Lpg1137, Cam^r^, Amp^r^ | This study |
| 362 pCS Cherry-*lpg1145* | Encoding mCherry-tagged Lpg1145, Cam^r^, Amp^r^ | This study |
| 362 pCS Cherry-*lpg1148* | Encoding mCherry-tagged Lpg1148, Cam^r^, Amp^r^ | This study |
| 362 pCS Cherry-*lpg1154* | Encoding mCherry-tagged Lpg1154, Cam^r^, Amp^r^ | This study |
| 362 pCS Cherry-*lpg1158* | Encoding mCherry-tagged Lpg1158, Cam^r^, Amp^r^ | This study |
| 362 pCS Cherry-*lpg1166* | Encoding mCherry-tagged Lpg1166, Cam^r^, Amp^r^ | This study |
| 362 pCS Cherry-*lpg1183* | Encoding mCherry-tagged Lpg1183, Cam^r^, Amp^r^ | This study |
| 362 pCS Cherry-*lpg1227* | Encoding mCherry-tagged Lpg1227, Cam^r^, Amp^r^ | This study |
| 362 pCS Cherry-*lpg1290* | Encoding mCherry-tagged Lpg1290, Cam^r^, Amp^r^ | This study |
| 362 pCS Cherry-*lpg1355* | Encoding mCherry-tagged Lpg1355, Cam^r^, Amp^r^ | This study |
| 362 pCS Cherry-*lpg1368* | Encoding mCherry-tagged Lpg1368, Cam^r^, Amp^r^ | This study |
| 362 pCS Cherry-*lpg1408* | Encoding mCherry-tagged Lpg1408, Cam^r^, Amp^r^ | This study |
| 362 pCS Cherry-*lpg1426* | Encoding mCherry-tagged Lpg1426, Cam^r^, Amp^r^ | This study |
| 362 pCS Cherry-*lpg1496* | Encoding mCherry-tagged Lpg1496, Cam^r^, Amp^r^ | This study |
| 362 pCS Cherry-*lpg1602* | Encoding mCherry-tagged Lpg1602, Cam^r^, Amp^r^ | This study |
| 362 pCS Cherry-*lpg1625* | Encoding mCherry-tagged Lpg1625, Cam^r^, Amp^r^ | This study |
| 362 pCS Cherry-*lpg1666* | Encoding mCherry-tagged Lpg1666, Cam^r^, Amp^r^ | This study |
| 362 pCS Cherry-*lpg1687* | Encoding mCherry-tagged Lpg1687, Cam^r^, Amp^r^ | This study |
| 362 pCS Cherry-*lpg1689* | Encoding mCherry-tagged Lpg1689, Cam^r^, Amp^r^ | This study |
| 362 pCS Cherry-*lpg1692* | Encoding mCherry-tagged Lpg1692, Cam^r^, Amp^r^ | This study |
| 362 pCS Cherry-*lpg1701* | Encoding mCherry-tagged Lpg1701, Cam^r^, Amp^r^ | This study |
| 362 pCS Cherry-*lpg1717* | Encoding mCherry-tagged Lpg1717, Cam^r^, Amp^r^ | This study |
| 362 pCS Cherry-*lpg1751* | Encoding mCherry-tagged Lpg1751, Cam^r^, Amp^r^ | This study |
| 362 pCS Cherry-*lpg1798* | Encoding mCherry-tagged Lpg1798, Cam^r^, Amp^r^ | This study |
| 362 pCS Cherry-*lpg1851* | Encoding mCherry-tagged Lpg1851, Cam^r^, Amp^r^ | This study |
| 362 pCS Cherry-*lpg1884* | Encoding mCherry-tagged Lpg1884, Cam^r^, Amp^r^ | This study |
| 362 pCS Cherry-*lpg1888* | Encoding mCherry-tagged Lpg1888, Cam^r^, Amp^r^ | This study |
| 362 pCS Cherry-*lpg1890* | Encoding mCherry-tagged Lpg1890, Cam^r^, Amp^r^ | This study |
| 362 pCS Cherry-*lpg1895* | Encoding mCherry-tagged Lpg1895, Cam^r^, Amp^r^ | This study |
| 362 pCS Cherry-*lpg1947* | Encoding mCherry-tagged Lpg1947, Cam^r^, Amp^r^ | This study |
| 362 pCS Cherry-*lpg1949* | Encoding mCherry-tagged Lpg1949, Cam^r^, Amp^r^ | This study |
| 362 pCS Cherry-*lpg1953* | Encoding mCherry-tagged Lpg1953, Cam^r^, Amp^r^ | This study |
| 362 pCS Cherry-*lpg1958* | Encoding mCherry-tagged Lpg1958, Cam^r^, Amp^r^ | This study |
| 362 pCS Cherry-*lpg1960* | Encoding mCherry-tagged Lpg1960, Cam^r^, Amp^r^ | This study |
| 362 pCS Cherry-*lpg1961* | Encoding mCherry-tagged Lpg1961, Cam^r^, Amp^r^ | This study |
| 362 pCS Cherry-*lpg1963* | Encoding mCherry-tagged Lpg1963, Cam^r^, Amp^r^ | This study |
| 362 pCS Cherry-*lpg1964* | Encoding mCherry-tagged Lpg1964, Cam^r^, Amp^r^ | This study |
| 362 pCS Cherry-*lpg1965* | Encoding mCherry-tagged Lpg1965, Cam^r^, Amp^r^ | This study |
| 362 pCS Cherry-*lpg1966* | Encoding mCherry-tagged Lpg1966, Cam^r^, Amp^r^ | This study |
| 362 pCS Cherry-*lpg1972* | Encoding mCherry-tagged Lpg1972, Cam^r^, Amp^r^ | This study |
| 362 pCS Cherry-*lpg1982* | Encoding mCherry-tagged Lpg1982, Cam^r^, Amp^r^ | This study |
| 362 pCS Cherry-*lpg2048* | Encoding mCherry-tagged Lpg2048, Cam^r^, Amp^r^ | This study |
| 362 pCS Cherry-*lpg2128* | Encoding mCherry-tagged Lpg2128, Cam^r^, Amp^r^ | This study |
| 362 pCS Cherry-*lpg2143* | Encoding mCherry-tagged Lpg2143, Cam^r^, Amp^r^ | This study |
| 362 pCS Cherry-*lpg2148* | Encoding mCherry-tagged Lpg2148, Cam^r^, Amp^r^ | This study |
| 362 pCS Cherry-*lpg2153* | Encoding mCherry-tagged Lpg2153, Cam^r^, Amp^r^ | This study |
| 362 pCS Cherry-*lpg2155* | Encoding mCherry-tagged Lpg2155, Cam^r^, Amp^r^ | This study |
| 362 pCS Cherry-*lpg2166* | Encoding mCherry-tagged Lpg2166, Cam^r^, Amp^r^ | This study |
| 362 pCS Cherry-*lpg2199* | Encoding mCherry-tagged Lpg2199, Cam^r^, Amp^r^ | This study |
| 362 pCS Cherry-*lpg2244* | Encoding mCherry-tagged Lpg2244, Cam^r^, Amp^r^ | This study |
| 362 pCS Cherry-*lpg2324* | Encoding mCherry-tagged Lpg2324, Cam^r^, Amp^r^ | This study |
| 362 pCS Cherry-*lpg2328* | Encoding mCherry-tagged Lpg2328, Cam^r^, Amp^r^ | This study |
| 362 pCS Cherry-*lpg2351* | Encoding mCherry-tagged Lpg2351, Cam^r^, Amp^r^ | This study |
| 362 pCS Cherry-*lpg2385* | Encoding mCherry-tagged Lpg2385, Cam^r^, Amp^r^ | This study |
| 362 pCS Cherry-*lpg2406* | Encoding mCherry-tagged Lpg2406, Cam^r^, Amp^r^ | This study |
| 362 pCS Cherry-*lpg2411* | Encoding mCherry-tagged Lpg2411, Cam^r^, Amp^r^ | This study |
| 362 pCS Cherry-*lpg2422* | Encoding mCherry-tagged Lpg2422, Cam^r^, Amp^r^ | This study |
| 362 pCS Cherry-*lpg2452* | Encoding mCherry-tagged Lpg2452, Cam^r^, Amp^r^ | This study |
| 362 pCS Cherry-*lpg2456* | Encoding mCherry-tagged Lpg2456, Cam^r^, Amp^r^ | This study |
| 362 pCS Cherry-*lpg2498* | Encoding mCherry-tagged Lpg2498, Cam^r^, Amp^r^ | This study |
| 362 pCS Cherry-*lpg2519* | Encoding mCherry-tagged Lpg2519, Cam^r^, Amp^r^ | This study |
| 362 pCS Cherry-*lpg2529* | Encoding mCherry-tagged Lpg2529, Cam^r^, Amp^r^ | This study |
| 362 pCS Cherry-*lpg2539* | Encoding mCherry-tagged Lpg2539, Cam^r^, Amp^r^ | This study |
| 362 pCS Cherry-*lpg2546* | Encoding mCherry-tagged Lpg2546, Cam^r^, Amp^r^ | This study |
| 362 pCS Cherry-*lpg2587* | Encoding mCherry-tagged Lpg2587, Cam^r^, Amp^r^ | This study |
| 362 pCS Cherry-*lpg2745* | Encoding mCherry-tagged Lpg2745, Cam^r^, Amp^r^ | This study |
| 362 pCS Cherry-*lpg2759* | Encoding mCherry-tagged Lpg2759, Cam^r^, Amp^r^ | This study |
| 362 pCS Cherry-*lpg2793* | Encoding mCherry-tagged Lpg2793, Cam^r^, Amp^r^ | This study |
| 362 pCS Cherry-*lpg2831* | Encoding mCherry-tagged Lpg2831, Cam^r^, Amp^r^ | This study |
| 362 pCS Cherry-*lpg2871* | Encoding mCherry-tagged Lpg2871, Cam^r^, Amp^r^ | This study |
| 362 pCS Cherry-*lpg2911* | Encoding mCherry-tagged Lpg2911, Cam^r^, Amp^r^ | This study |

**Table S3. Oligonucleotides used in this study.**

| **Name** | **Sequence 5’ to 3’ direction** |
| --- | --- |
| Fwd-attB1-*lpg0696*_407-570_ | GGGGACAAGTTTGTACAAAAAAGCAGGCTTATTAGATGAGCTCATACG |
| Rev-attB2-*lpg0696*_407-570_ | GGGGACCACTTTGTACAAGAAAGCTGGGTTCTATTTCAATTCATTTTTATTTTCTATTTC |
| Fwd-attB1-*lpg1798*_221-336_ | GGGGACAAGTTTGTACAAAAAAGCAGGCTTAGAAAGGCATGCCGATAAATT |
| Rev-attB2-*lpg1798*_221-336_ | GGGGACCACTTTGTACAAGAAAGCTGGGTTTTAAGAGCGCGATAT |
| Fwd-attB1-*lpg1963*_361-499_ | GGGGACAAGTTTGTACAAAAAAGCAGGCTTAAAAGACCAAAAAGAGTTTTCTAAATT |
| Rev-attB2-*lpg1963*_361-499_ | GGGGACCACTTTGTACAAGAAAGCTGGGTTTTATCTCTTTTGAACCTG |
| Fwd-attB1-*lpg2153*_902-1121_ | GGGGACAAGTTTGTACAAAAAAGCAGGCTTACGACATGAAAGCGG |
| Rev-attB2-*lpg2153*_902-1121_ | GGGGACCACTTTGTACAAGAAAGCTGGGTTTTAAATTGGAGTAGTGGT |
| Fwd-attB1-*lpg2385*_289-447_ | GGGGACAAGTTTGTACAAAAAAGCAGGCTTATCAGATGAAAAAATTGACGTAGT |
| Rev-attB2-*lpg2385*_289-447_ | GGGGACCACTTTGTACAAGAAAGCTGGGTTTTACGTTTTTGCTAC |
| Fwd-attB1-*lpg2546*_278-464_ | GGGGACAAGTTTGTACAAAAAAGCAGGCTTACGATTTGGAAAAGATGA |
| Rev-attB2-*lpg2546*_278-464_ | GGGGACCACTTTGTACAAGAAAGCTGGGTTTTAAGATGAATGCTC |
| Fwd-attB1-*lpg2587*_391-516_ | GGGGACAAGTTTGTACAAAAAAGCAGGCTTATCCATGGTCAAAACCGATAA |
| Rev-attB2-*lpg2587*_391-516_ | GGGGACCACTTTGTACAAGAAAGCTGGGTTTTAGTCTGTTGTTTTTGCCAG |
